# Putative EGF ligand and receptor of *Echinococcus multilocularis* that are critical for parasite development

**DOI:** 10.1101/2024.12.20.629808

**Authors:** Akito Koike, Katia Cailliau, Jérôme Vicogne, Frank Becker, Colette Dissous, Stefan Hannus, Klaus Brehm

## Abstract

The neglected zoonosis alveolar echinococcosis (AE) is caused by infiltrative growth of the metacestode larval stage of the cestode *Echinococcus multilocularis* within host organs. We previously demonstrated that metacestode growth depends on the mitotic activity of a population of parasite stem cells, called germinative cells, but it is not yet clear which molecular mechanisms govern *Echinococcus* stem cell dynamics such as cell-cycle progression, self-renewal and differentiation. Based on previous reports showing that epidermal growth factor (EGF) signalling contributes to *Echinococcus* stem cell regulation, we herein characterized three EGF receptors of the parasite and demonstrated by RNAi and inhibitor assays that one of these, EmER1, is crucial for the development of metacestode vesicles from parasite stem cells. We also showed that EmER1 serves as a target for afatinib, an EGF receptor inhibitor with profound anti-parasitic activities *in vitro* and *in vivo*. By bioinformatic analyses and membrane-bound yeast two-hybrid assays, we identified a parasite-derived, neuregulin-like cognate ligand for EmER1, EmNRG, the expression of which is strongly upregulated in metacestode vesicles during clonal expansion of germinative cells. Furthermore, we demonstrate that RNAi knockdown of the EmNRG encoding gene drastically affects the ability of germinative cells to produce metacestode vesicles. We propose that EmNRG and EmER1 form a cognate ligand-receptor system utilized by *E. multilocularis* to regulate asymmetric versus symmetric division decisions of stem cells. These data are relevant for further studies into *Echinococcus* stem cell dynamics and for the development of EGF signalling-based anti-infectives against echinococcosis.

## Introduction

Alveolar echinococcosis (AE) is a potentially lethal zoonosis caused by the larval stage of the fox-tapeworm *Echinococcus multilocularis*. Infection of the intermediate host (rodents, humans) is initiated by the oral uptake of infectious eggs that contain the parasite’s oncosphere stage (Thompson, 2017). Within the intestine, the oncosphere hatches from the egg penetrates the intestinal barrier, and gains access to the internal organs where, mostly within the liver, it undergoes a metamorphosis towards the metacestode stage (Brehm and Koziol, 2017; Thompson, 2017). The metacestode is basically a cyst-like structure surrounded by an acellular laminated layer, involved in host protection, and an inner cellular ‘germinal layer’ made up of few cell types such as stem-, muscle-, nerve-, and glycogen storing-cells as well as a syncytial tegument (Brehm and Koziol, 2017). In the case of *E. multilocularis*, the multivesicular metacestode tissue grows cancer-like infiltratively into the liver tissue and we previously demonstrated that parasite proliferation is exclusively driven by a stem cell population (also called ‘germinative cells’), which are the only mitotically active parasite cells and give rise to all differentiated parasite cells (Koziol et al., 2014). This study and recent work of our laboratory concerning the germinative cell transcriptome (Herz et al., 2024) also showed that the *Echinococcus* stem cell system displays several similarities (but also some notable differences) to the neoblast stem cells of free-living flatworms. It is assumed that the *Echinococcus* germinative cells are relatively resistant to the current drugs of choice against AE, albendazole and mebendazole, and that novel medication should primarily target the parasite’s stem cell system to achieve parasitocidal effects (Brehm and Koziol, 2014; Koziol and Brehm, 2015). Due to the importance of germinal cells for parasite persistence within the host, their intrinsic mechanisms regulating stem cell renewal and differentiation towards muscle-nerve-, and tegumental cells are of special medical and scientific interest.

One of the evolutionarily conserved signalling pathways that regulate differentiation of metazoan stem cells is the Epidermal Growth Factor (EGF) pathway, which involves secreted (and sometimes membrane bound) ligands of the EGF/neuregulin type and receptor tyrosine kinases of the EGF-receptor (EGFR) families (Perrimon et al., 2012; Marchionni, 2014). Upon binding of the EGF family ligand to cognate receptors, these usually undergo dimerization and cross-phosphorylation, eventually activating downstream acting signalling cascades such as the Erk-like mitogen activated protein kinase (MAPK) or the Akt-PI3K-mTOR pathways (Burgess, 2022). Importantly, previous work on the free-living flatworm species *Schmidtea mediterranea* revealed that neoblast self-renewal and differentiation is decisively regulated by EGF signalling (Lei et al., 2016). These authors demonstrated that *S. mediterranea* expresses one EGF ligand, and one corresponding receptor, which direct asymmetric division of neoblasts towards progeny cells with properties of differentiating cells and stem cells. In *Echinococcus*, previous work indicated the presence of at least one EGFR-like receptor, named EmER (Spiliotis et al., 2003), and putative downstream acting components belonging to Erk-like MAPK cascade pathways (Spiliotis et al., 2006; Gelmedin et al., 2010). Furthermore, Cheng et al. showed that germinative cell proliferation is stimulated by exogenous addition of human EGF and that metacestode growth is inhibited in the presence of potential EGFR inhibitors, such as afatinib, and inhibitors of the Erk-like MAPK cascade branch (Cheng et al., 2017). In a subsequent study, these authors also demonstrated the induction of apoptosis in germinative cells upon addition of EGFR inhibitors (Cheng et al., 2020), indicating that, like in planarians, EGF signalling also governs stem cell differentiation/self-renewal in cestodes.

Although previous analyses by Cheng et al. indicated that host-derived EGF can stimulate the parasite EGFR EmER *in vitro*, clear effects could only be observed at concentrations of 10 ng/ml and higher (Cheng et al., 2017), which exceeds physiological concentrations of EGF in human tissues (1-2 ng/ml; (Pinilla-Macua et al., 2017)). Even considering that the expression of EGF is induced during liver regeneration (Mullhaupt et al., 1994), it is therefore questionable whether concentrations of host EGF that are relevant for stimulating parasite development can be physiologically achieved during an infection by *E. multilocularis*. This, together with the fact that free-living flatworms express several EGF ligands (Barberan et al., 2016), prompted us to investigate whether *E. multilocularis* itself expresses such ligands for parasite-intrinsic regulation of stem cell proliferation. We herein report the identification of two *Echinococcus* EGF ligands and two additional EGFR-like receptors and show expression of EGF ligand and receptor genes in parasite stem cells as well as differentiated cells. We further demonstrate interaction between the *Echinococcus* ligand and EGF receptors and show that RNAi knockdown of the respective genes significantly affects parasite development. Finally, we demonstrate that the parasite’s EGF ligand is drastically induced during clonal expansion of germinative cells, and we identified additional inhibitors affecting the activity of *Echinococcus* EGF receptors. Our data are discussed in the context of *Echinococcus* stem cell regulation and drug development.

## Materials and Methods

### Ethics statement

*In vivo* propagation of parasite material was performed in Mongolian jirds (*Meriones unguiculatus*), which were raised and housed at the local animal facility of the Institute of Hygiene and Microbiology, University of Würzburg. This study was performed in strict accordance with German (*Deutsches Tierschutzgesetz,TierSchG*, version from Dec-9-2010) and European (European directive 2010/63/EU) regulations on the protection of animals. The protocol was approved by the Ethics Committee of the Government of Lower Franconia (Regierung von Unterfranken) under permit numbers 55.2–2531.01-61/13 and 55.2.2-2532-2- 1479-8.

For *Xenopus laevis*, experiments were performed according to the European Community Council guidelines (86/609/EEC), and the protocol was approved by the local institutional review board (Comité d’Ethique en Expérimentation Animale, Haut de France, France, G59- 00913).

### Organisms and culture methods

All inhibitor, RNAi, and *in situ* hybridization experiments were performed with two isolates of *E. multilocularis*, H95 from a naturally infected fox (*Vulpes vulpes*) (Jura et al., 1996) and GH09 from a naturally infected cynomolgus monkey (*Macaca fascicularis*) (Tappe et al., 2007). The isolates have been passaged in Mongolian jirds (*Meriones unguiculatus*) as described (Spiliotis et al., 2004; Spiliotis and Brehm, 2009). Preparation and use of conditioned media for axenic culture were described previously (Koike et al., 2022).

*Xenopus laevis* (wild type strain from the CRB-University of Rennes, France) were purchased from TEFOR Paris-Saclay Zootechnie aquatique (TPS-AQUA), France. They were housed in PHExMAR, University of Lille, at 18°C in the Xenoplus Housing System (Techniplast).

Yeast Two-Hybrid (Y2H) Gold strain from Clontech (*MATa, trp1-901, leu2-3, 112, ura3-52, his3-200, gal4Δ, gal80Δ, LYS2:: GAL1_UAS_–Gal1_TATA_–His3, GAL2_UAS_–Gal2_TATA_–Ade2, URA3::MEL1_UAS_–Mel1_TATA_, AUR1-C MEL1*) used in yeast two hybrid experiments was passaged once a month on conventional YPD plates and kept at 4℃. Dropout medium and plates were prepared with Difco^TM^ Yeast Nitrogen Base without Amino Acid (Becton Dickinson) and DO supplements (Clontech) following the manufacturer’s instructions.

### Anti-parasitic inhibitor assays

Providers of protein kinase inhibitors are listed in Table S1. Primary cells for cell viability assay and vesicle regeneration assay were isolated from *in vitro* cultivated metacestode vesicles using a previously established protocol (Spiliotis et al., 2010). Primary cell unit definition and experimental procedure for cell viability and regeneration assay were described previously (Koike et al., 2022). The respective experiments were performed in 3 technical replicates and 3 biological replicates. The number of newly regenerated vesicles was analyzed in principle with Kruskal-Wallis tests followed by Dunn’s multiple comparison tests in Graphpad Prism 10.1.2 (Graphpad software). The procedure of the mature vesicle assay is described by Koike et al. (Koike et al., 2022). In this experiment, 3 biological replicates were used. The percentage of the intact vesicles in the mature vesicle assay and that of cell viability were analyzed in principle with one-way-ANOVA followed by Dunnett’s multiple comparison tests in Graphpad Prism 10.1.2 (Graphpad software).

### Nucleic acid isolation, cloning, sequencing, and *in vitro* synthesis of cRNA

RNA isolation from HEK293T cells was performed using a Trizol-based method as described (Hemer et al., 2014). From total RNA, cDNA was synthesized with oligo(dT)20 primer and Superscript IV reverse transcriptase (Invitrogen) following the manufacturer’s instruction. This cDNA was used as the PCR template for the construct of HsEGF. For the constructs of HER1, pMK-RQ-ErbB1 was used as a PCR template. For other constructs of *Echinococcus*, pJG4-5 based yeast-two-hybrid library (Hubert et al., 2004) were used as PCR templates. PCR primers were purchased from Merck (Germany) and are listed in supplementary Table S2. Purified DNA fragments were cloned into linearized plasmids and sequenced by Sanger Sequencing (Microsynth seqlab). Sequences of newly analyzed genes in this work have been deposited in the GenBank database and accession numbers are listed in Table S3. Prior to synthesis of capped mRNA (cRNA), pSecTag2Hygro-based plasmids (EmER1, EmER2, EmER3) and pcDNA3.1-SER (Vicogne et al., 2004) were linearized by Pme I and pOBER plasmid including HER1 (Opresko and Wiley, 1990) was linearized by Not I. By using these linearized vectors as templates, cRNA was synthesized from these vectors as templates using the T7 mMessage mMachine Kit (Ambion). After purification and quantification, the size of the RNA fragment and the lack of abortive transcripts were confirmed through gel electrophoresis.

### Expression of EGFRs in *Xenopus* oocytes

Oocytes of *Xenopus laevis* were surgically removed from animals and prepared for the microinjection, as described (Vicogne et al., 2004). Oocyte viability was evaluated by counting GVBD+ (germinal vesicle breakdown) oocytes without microinjection, after progesterone treatment. 60 ng of *emer1*, *emer2*, *emer3*, *SER*, and human *HER1* cRNA were injected into oocytes. 8 or 24 h after microinjection of cRNA, the oocytes were stimulated by 50 nM recombinant human EGF or progesterone. GVBD was evaluated 15 h after progesterone/EGF stimulation.

For immunoprecipitation and western blot analysis, oocytes were lysed 15h after expression in buffer PY (20 mM Tris-HCl pH 7.4, 50 mM NaCl, 5 mM EDTA, Triton X-100 1%, 1 mg/ml bovine serum albumin, 10 µg/ml leupeptin, 10 µg/ml aprotinin, 10 µg/ml soybean trypsin inhibitor, 10 µg/ml benzamidine, 1 mM PMSF, 1 mM sodium vanadate) and centrifuged at 4°C for 15 min at 10,000 g. Supernatants were incubated for 90 min at 4°C with anti-myc antibodies (Santa Cruz Biotechnology,1/200) in the presence of protein A-sepharose beads (1/20 µl, Sigma). Immune complexes eluted were separated on 4-20% SDS PAGE gels (mini protean TGX, BioRad) for 1 h at 200 V. Transfer was performed on nitrocellulose (Amersham Hybond) membrane at 100 V for 1 h using buffer B (0.32% TRIS; 1.8% glycine; 20% methanol). Membranes were cut according to the standard molecular weight of proteins, saturated, and incubated overnight at 4°C with antibodies raised against phosphotyrosine PY20 (Santa Cruz Biotechnology, 1/1200), before treatment with a ready-to-use stripping buffer (Clinisciences) and incubation with anti-myc antibodies (Santa Cruz Biotechnology, 1/1000). Membranes were revealed with Select ECL detection system (Amersham).

To evaluate the effect of inhibitors, oocytes with or without cRNA microinjection were treated with various concentrations of inhibitors, dacomitinib, afatinib, and osimertinib for 30 min before EGF/progesterone stimulation. All experiments were performed using 2-3 technical replicates. Percentages of GVBD-positive oocytes were statistically analyzed with one-way ANOVA with Dunnett’s multiple comparison tests in Graphpad Prism 10.1.2 (Graphpad software).

### Bioinformatic analysis

Bioinformatic analysis was essentially performed as described (Koike et al., 2022). Briefly, BLASTP searches against protein databases of *E. multilocularis*, *Schistosoma mansoni,* and *Schmidtea mediterranea* were performed on Wormbase ParaSite (Howe et al., 2017) using the amino acid sequences of human EGF receptors and EGF ligands as queries. The KEGG database at Genomenet (https://www.genome.jp/) was used for the reciprocal BLASTP analysis. Gene predictions based on the *E. multilocularis* genome sequence (Tsai et al., 2013) available in Wormbase ParaSite, were verified or corrected using the Integrated Genome Viewer (Thorvaldsdóttir et al., 2013), and previously generated transcriptome data (Tsai et al., 2013; Herz et al., 2024). For domain detection, SMART (Letunic et al., 2021) was used. Multiple sequence alignments were performed using CLUSTALW 2.1 (Thompson et al., 1994).

### *In situ* hybridization and 5-ethynyl-2’-deoxyuridine (EdU) labeling

Fragments of the *emnrg* cDNA were amplified from a pJG4-5-based yeast-two-hybrid library (Hubert et al., 2004) and cloned into pJET1.2 with CloneJET PCR Cloning Kit (Thermo Fisher Scientific). Primers used for cloning are listed in Table S2. Digoxygenin (DIG)-labeled probes were synthesized from this pJet 1.2-based plasmid, purified, and quantified as described (Koike et al., 2022). *In vitro* cultivated metacestode vesicles for WISH were treated with 40mM hydroxyurea (HU) for 7 days as described (Koziol et al., 2014) and recovered for 0, 3, 6, or 10 days. All vesicles were labeled with 50μM 5-ethynyl-2’-deoxyuridine (EdU) *in vitro* at 37℃ for 5 h prior to fixation by 4% paraformaldehyde. Whole-mount *in situ* hybridization (WISH) followed by fluorescent detection of EdU with Alexa Flour 555 was carried out as previously described (Koziol et al., 2014). Fluorescent specimens were imaged with a Nikon Eclipse Ti2E confocal microscope and processed with Fiji/ImageJ as described (Koike et al., 2022). One-way ANOVA followed by Dunnett’s multiple comparison test was used in Graphpad Prism 10.1.2 (Graphpad software) to compare the number of cells with each signal.

### Membrane-anchored ligand and receptor yeast two-hybrid (MALAR-Y2H) analyses

The previously established MALAR-Y2H system (Li et al., 2016) was used for assessing ligand-receptor interactions, with plasmids pGAD_SP-WBP1_cloning_ linker_TMP_Cub_GAL4 (for ligands), and pGBKT7_SP_OST1_cloning_NubG (for receptors). cDNA fragments encoding sequences downstream of the signal peptide and upstream of the transmembrane domain of the *Echinococcus* ligand cDNA were amplified from pJC4-5 based plasmid library (Hubert et al., 2004), and integrated into *SmaI*-digested pGAD_SP-WBP1_cloning_linker_TMP_Cub_GAL4. For the human EGF construct, cDNA synthesized from HEK293T cells was used for the PCR template. Similarly, fragments between signal peptide and juxtamembrane region of the receptor cDNA were amplified and integrated into *SmaI*-digested pGBKT7_SP_OST1_cloning_NubG. Primers used for cloning are listed in Table S2. The *Saccharomyces cerevisiae* Y2HGold strain (Clontech) was transformed by these plasmids using the protocol of Tripp et al (Tripp et al., 2013). Transformants were initially inoculated onto Leu-/Trp-double dropout plates, and later transferred onto Leu-/Trp-/His-triple dropout plate and Leu-/Trp-/Ade-/His-quadruple dropout plates with three densities, as described (Koike et al., 2022). Growth level was quantified by Fiji/ImageJ using the protocol of Petropavlovskiy et. al. (Petropavlovskiy et al., 2020). The quantified level of growth on quadruple dropout plates was statistically analyzed using one-way-ANOVA followed by Tukey’s multiple comparison tests in Graphpad Prism 10.1.2 (Graphpad software).

### RNA interference

RNA interference (RNAi) was performed using a slightly modified protocol of Spiliotis et al. (Spiliotis et al., 2010). siPOOLs (mixture of 30 siRNA fragments) were designed and synthesized by siTOOLs Biotech (Planegg, Germany). Other siRNAs were designed with siDirect version 2.0 (Naito et al., 2009) and purchased from Sigma-Aldrich. As a negative control, randomized siRNA fragments from siTOOLs and siRNA against eGFP from Sigma-Aldrich were used respectively. The sequences of the siRNAs from Sigma-Aldrich are listed in Table S4. *E. multilocularis* primary cells for RNAi experiments were isolated with the same procedure as those for cell viability assay and vesicle regeneration assay. 150 U of primary cells isolated from mature metacestode vesicles were dispensed into an electroporation cuvette (1mm Gap) from BTX with 90 µl of electroporation buffer (siPort) including 200pmol siRNA fragments and electroporated with time constant protocol at 200 V for 0.5 ms with the electroporator GenePulser Xcell (BioRad). The electroporated primary cells were incubated for 12 min at 37℃ and centrifuged at room temperature. After removal of the supernatant, cells were resuspended by the A6/B4 condition medium and dispensed into a 96-well plate (150 U/well). Primary cells were then cultured for 21 days, and half of the conditioned medium was exchanged twice a week. Images were taken under an optical microscope (Nikon eclipse Ts2-FL) and the number of regenerated vesicles was counted. Three technical replicates and three biological replicates were prepared for counting regenerated vesicles. With only three technical replicates, cell viability was measured with Cell Titer Glo (Promega) after 21 days of incubation.

## Results

### Cloning and characterization of *Echinococcus* EGF receptors

Prompted by the importance of stem cells in metacestode proliferation (Koziol et al., 2014) and the role of EGF signalling in flatworm stem cell biology (Lei et al., 2016), we were interested in characterizing the full EGF receptor complement of *E. multilocularis*. One receptor of this family, EmER, had already been characterized previously and displayed an EGFR typical structure of an intracellular tyrosine kinase domain as well as two extracellular receptor L domains together with several furin-like domains (Spiliotis et al., 2003). The combination of intracellular tyrosine kinase domains with extracellular receptor L domains is a hallmark of the EGF and insulin receptor families (Ward et al., 2007) and when we inspected the previously characterized *E. multilocularis* genome sequence (Tsai et al., 2013), only five gene models were predicted to encode proteins of this composition. Two of those (EmuJ_000962900, EmuJ_000981300) encoded members of the insulin receptor family which we had previously characterized (Konrad et al., 2003; Hemer et al., 2014), one referred to EmER (EmuJ_000075800), and two others (EmuJ_000617300, EmuJ_000969600) were annotated as receptor tyrosine kinases. BLASTP analyses then revealed that the encoded proteins displayed the highest similarities to EGF receptors of different metazoan origin. We thus concluded that, based on genome sequence predictions, EmuJ_000617300 and EmuJ_000969600 represented two additional genes for EGF receptor family members, which is in line with the results of a previous study that had been conducted on the evolution of EGF receptors in metazoans (Barberan et al., 2016). To ensure that all respective genes had been predicted during genome annotation, we also carried out TBLASTN analyses against the entire genome sequence using the tyrosine kinase domains and the receptor L regions of EmuJ_000075800, EmuJ_000617300, and EmuJ_000969600 as queries but did not find additional genes. Hence, in contrast to planarians, which express 6 different EGF receptor genes (Barberan et al., 2016), the *E. multilocularis* genome only encodes 3 of these receptors. Informed by the *E. multilocularis* genome sequence and recently generated transcriptome analyses (Herz et al., 2024), we then generated primers to full-length clone the remaining *Echinococcus* EGF receptors and deposited the respective cDNA sequences in Genbank (Table S3). It should be noted that in the literature there is some confusion concerning the designation of *Echinococcus* EGF receptors. Whilst the first cloned EGF receptor gene was named *emer* (Spiliotis et al., 2003), referring to model EmuJ_000075800, Barberán et al. designated this gene *EGFR-2* (Barberan et al., 2016). These authors also allocated *EGFR-1* to EmuJ_000617300 and *EGFR-3* to EmuJ_000969600, whereas we in one of our first genome analyses named these genes *emerb* and *emerc* (Brehm and Koziol, 2017). To avoid confusion and to harmonize the designation of *Echinococcus* EGF receptor gene with those of insulin-(Hemer et al., 2014), fibroblast growth factor-(Förster et al., 2018), and transforming growth factor receptor genes (Kaethner et al., 2023), we finally decided to dub them *emer1* (EmuJ_000075800), *emer2* (EmuJ_000617300), and *emer3* (EmuJ_000969600), encoding the proteins EmER1, EmER2, and EmER3, respectively. It should also be noted that while the cDNA sequences encoding EmER1 and EmER3 were correctly predicted during annotation, the predicted cDNA for EmER2 was too short. The true cDNA encoded a protein with 31 additional amino acids at the N-terminus, which also contained an N-terminal signal peptide.

As shown in Figure 1A, the three *Echinococcus* EGF receptors contained predicted tyrosine kinase domains in the intracytoplasmic region as well as export-directing signal peptides, and two receptor L domains. In the case of EmER2, the SMART algorithm (Letunic et al., 2021) did not identify transmembrane helices. However, hydrophilicity plotting (Kyte and Doolittle, 1982) identified a region of high hydrophobicity between amino acids 979 and 1002 of EmER2 and we propose that this is the protein’s transmembrane region. We also carried out sequence homology analyses within the tyrosine kinase domains of all three *Echinococcus* EGF receptors and, as shown in Figure 1B, the proteins contained all residues highly conserved among EGF receptor-like tyrosine kinases (Hanks et al., 1988) at the corresponding positions, indicating that they are active tyrosine kinases. Especially, all of EmEGFR1-3 have catalytic lysine (Figure 1B), and at the position of asparagine 834 of HER3, which makes it catalytically impaired (Rubin and Yarden, 2001; Littlefield et al., 2014), all of them have aspartate (catalytic loop in Fig1B, Figure S5A).

**Figure 1:**
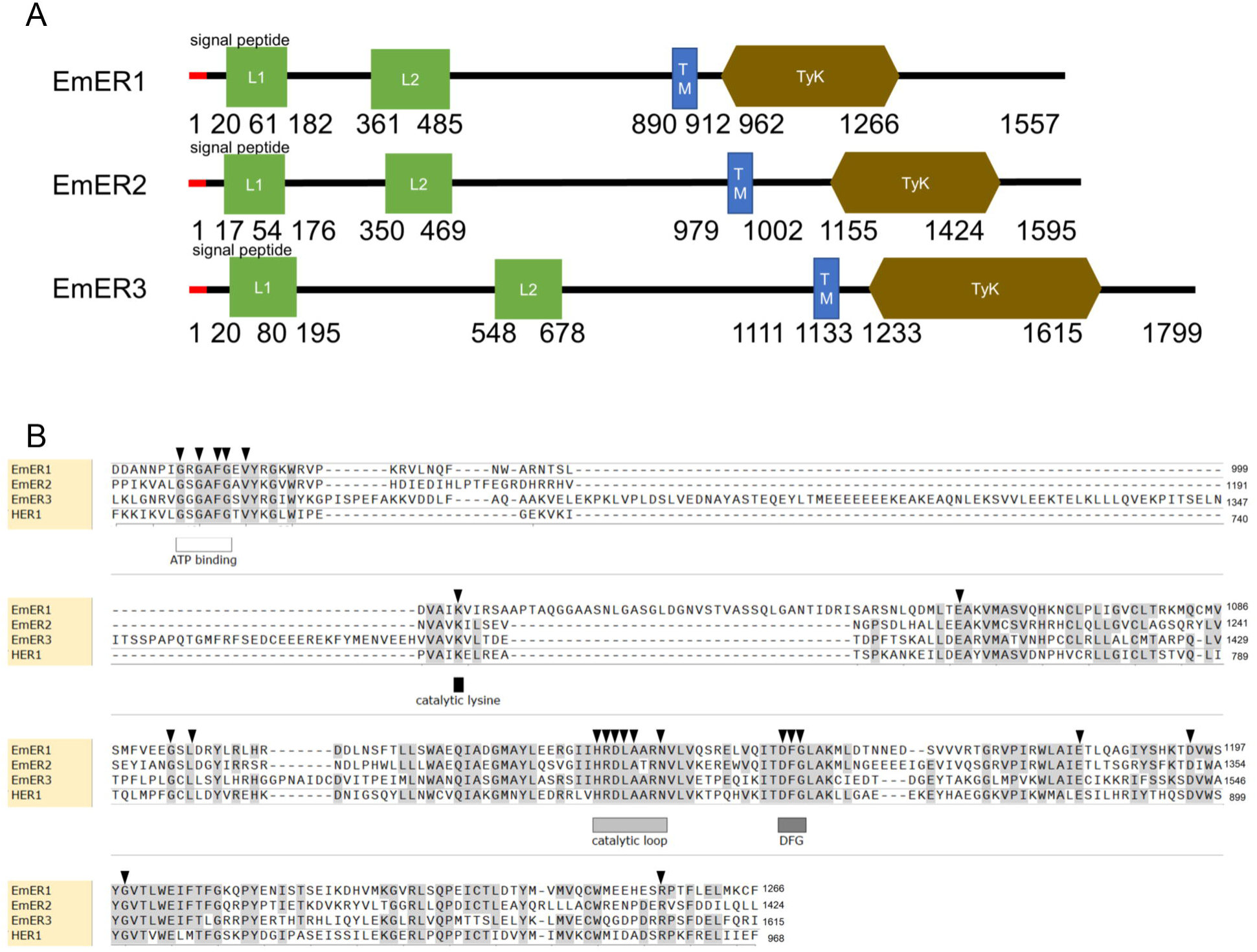
Domain structure and homologies of *Echinococcus* EGF receptors. (A) Domain structure of *Echinococcus* receptors EmER1, EmER2, and EmER3. Shown are the positions of the following domains: tyrosine kinase (TyK, brown), transmembrane (TM, blue), and receptor-L-domain (L1/L2, green), as well as N-terminal signal peptides (red). The number below indicates the position and length of the domains. (B) Amino acid sequence comparison of the tyrosine kinase domains of *Echinococcus* EGF receptors and HER1 (as indicated). Black triangles on the alignment indicate highly conserved residues among EGF receptor-like tyrosine kinases (Hanks et al., 1988). The locations of ATP-binding, DFG motif, catalytic lysin, and catalytic loop are shown.

### Expression patterns of *Echinococcus* EGF receptor genes

We were interested in the expression profiles of *emer1*, *emer2*, and *emer3* in *Echinococcus* developmental stages and inspected transcriptome datasets which had previously been generated for protoscoleces (before and after activation by low pH/pepsin treatment) and adult worms (Tsai et al., 2013) as well as metacestode vesicles (before and after treatment with hydroxyurea, thus eliminating stem cells) and parasite primary cell cultures at different stages of development (Herz et al., 2024). As shown in Figure 2, all three genes were well expressed in metacestode vesicles in which *emer2* showed the lowest level of expression. Likewise, all three genes were well expressed in protoscoleces, again with the lowest expression levels for *emer2*. None of the EGF receptor encoding genes displayed significantly lower TPM levels after treatment of metacestode vesicles with hydroxyurea, which we recently showed to be characteristic of stem cell-associated *Echinococcus* genes (Herz et al., 2024), indicating that they are predominantly expressed in differentiated (or differentiating) parasite cells. Rather low overall expression levels were observed for adult worms and when we inspected a previously generated transcriptome dataset for oncospheres (Huang et al., 2016), significant levels of expression were only observed for *emer1* and *emer3*. Hence, over the entire parasite life cycle *emer1* and *emer3* displayed prominent expression whereas *emer2* appears to be less well expressed.

**Figure 2:**
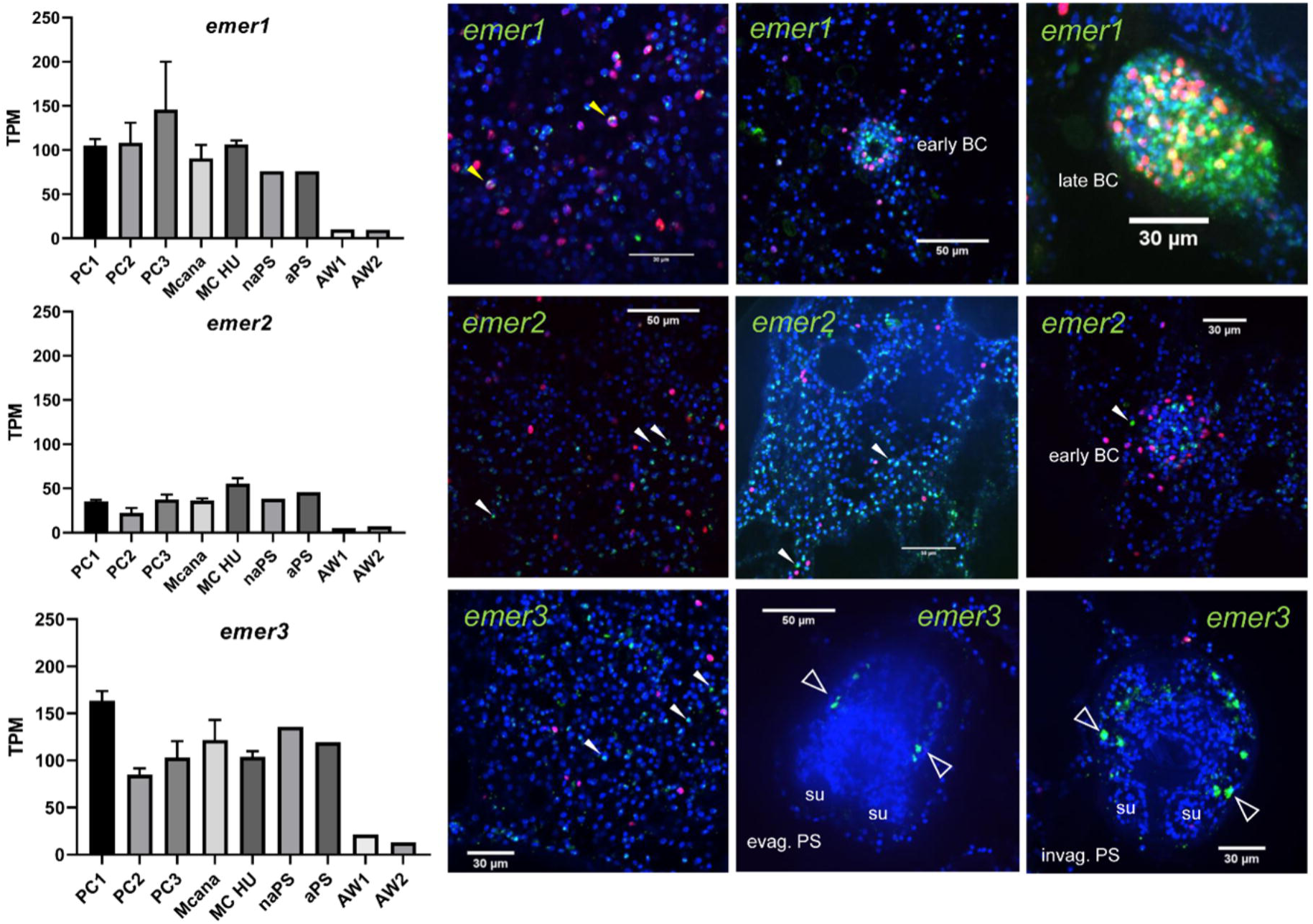
Expression of EGF receptor genes in *Echinococcus* larvae. Left panel: Expression values (in TPM) of *Echinococcus* EGF receptor encoding genes *emer1*, *emer2*, and *emer3* in primary cell cultures after 2 (PC1), 7-11 (PC2), and 16-22 days (PC3), *in vitro* cultivated metacestode vesicles before (MCana) or after (MC HU) treatment with hydroxyurea, non-activated (naPS) and activated (aPS) protoscoleces as well as pre-gravid (AW1) and gravid (AW2) adult worms (all according to Herz et al., 2024 and Tsai et al., 2013). Right panels: WISH for *emer1*, *emer2*, and *emer3* (as indicated) on metacestode vesicles at different phases of development and protoscoleces. Shown are for *emer1*: (from left to right): germinal layer, germinal layer with early brood capsule, and late brood capsule; for *emer2*: germinal layer (twice), germinal layer with early brood capsule; for *emer3*: germinal layer, evaginated protoscolex, invaginated protoscolex. Yellow triangles indicate WISH+/EdU+ cells, white triangles indicate cells only positive for WISH. Open triangles indicate *emer3*+ cells laterally positioned in the protoscolex. Shown are merge images (single confocal slices) of all three channels (blue, DAPI, nuclei; red, EdU, S-phase stem cells; green, WISH). BC, brood capsule; SU, sucker.

To further characterize the expression patterns of *Echinococcus* EGF receptor genes, we performed *in situ* hybridization on *in vitro* cultivated metacestode vesicles combined with EdU staining for labeling S-phase stem cells. As shown in Figure 2, we obtained numerous signals for all three genes that were dispersed over the metacestode germinal layer, but only very few cells with positive *in situ* hybridization signal also stained positive for EdU (less than 1% of all *in situ* hybridization positive cells). This indicated that the *Echinococcus* EGFR encoding genes are predominantly expressed in post-mitotic cells, which are either terminally differentiated or in a transition state. In the case of *emer1* and *emer2*, we also obtained strong signals in early brood capsules and, particularly for *emer1*, in late brood capsules where multiple EdU+ cells are found and protoscoleces are formed (Figure 2). In the case of *emer3*, we also detected a striking pattern of gene expression in lateral cells of the protoscolex (Figure 2), indicating that it might be involved in protoscolex pattern formation.

Taken together, our analyses showed that all three *Echinococcus* EGFR encoding genes are well expressed in the metacestode stage in post-mitotic cells and that all three genes are probably also involved in the formation of protoscoleces.

### *emer1* is necessary for metacestode formation

We had previously introduced a method to knock down the expression of *Echinococcus* genes in primary cell cultures using RNA interference (Spiliotis et al., 2010). Since all three EGFR encoding genes were well expressed in *Echinococcus* primary cells (Figure 2), we decided to investigate their role in metacestode formation using RNAi and combined our RNAi protocol with siPOOL technology (Hannus et al., 2014) to minimize off-target effects. We achieved gene knockdown to approximately 60% in all three cases three days after RNAi application. We then measured the formation of mature metacestode vesicles from these primary cell cultures and observed no development of vesicles after 21 days in the case of *emer1* (RNAi), whereas *emer2* (RNAi) showed no effect and in the case of *emer3* (RNAi) we even detected stimulation of vesicle development (Figure 3A). To further elucidate the cellular basis of reduced vesicle formation in the case of *emer1* (RNAi), we also measured the viability of primary cell cultures according to a previously established protocol (Koike et al., 2022). In these experiments, we observed significantly reduced cell viability after *emer1* (RNAi), whereas in the case of *emer2* and *emer3* overall viability was unaffected (Figure 3B). Taken together, our RNAi knockdown experiments indicated that *emer1* is necessary for metacestode vesicle formation and that reduced expression of this receptor affects *Echinococcus* cell viability.

**Figure 3:**
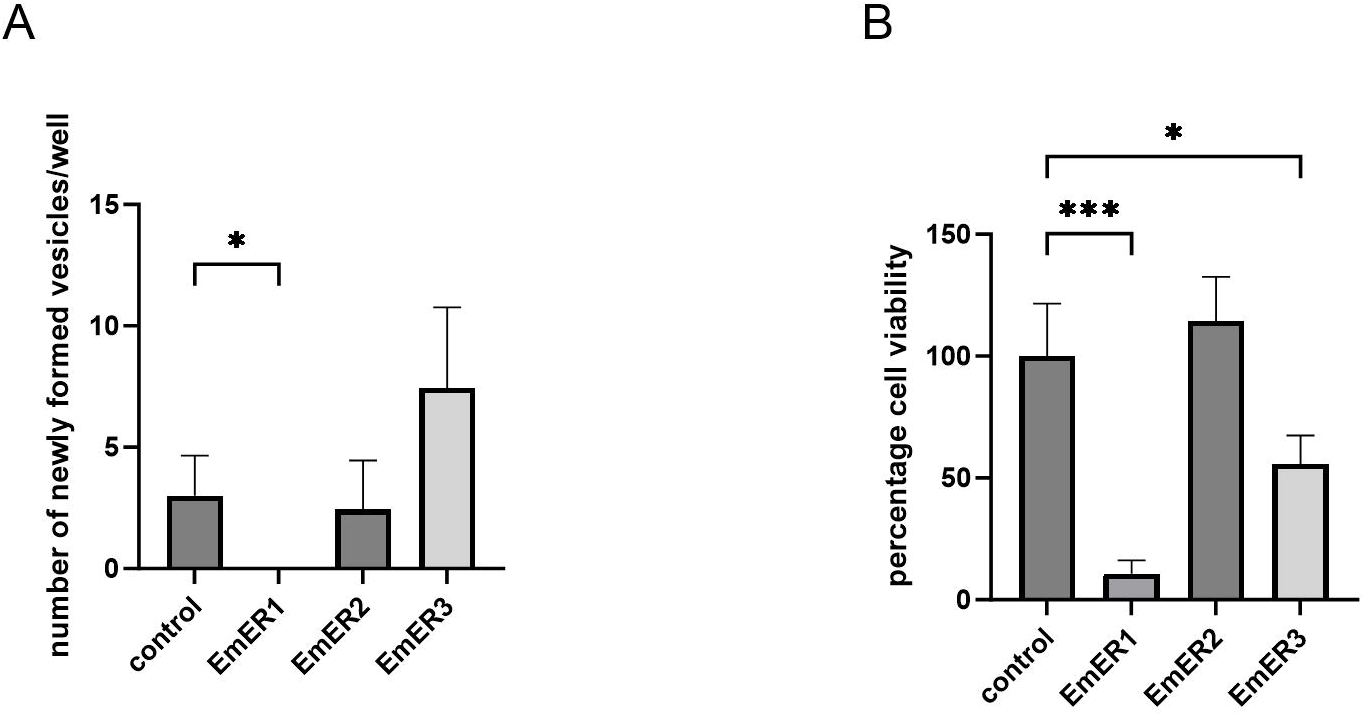
EmER1 is necessary for metacestode development. Displayed are the results of RNAi knockdown of genes encoding EmER1, EmER2, and EmER3 (as indicated). (A) Formation of mature metacestode vesicles from primary cell cultures after 21 days. The numbers of regenerated vesicles were analyzed by Kruscal-Wallis tests followed by Dunn’s tests. P values less than 0.0332 was summarized with *. (B) Cell viability of primary cell cultures after 3 days. Experiments have been performed as three biological triplicates. The normalized cell viability was analyzed by one-way ANOVA followed by Dunnet’s test. The error bar represents the standard deviation. P values less than 0.0332, and 0.0002 are summarized with * and ***.

### Afatinib targets EmER1

In a previous study, Cheng et al. demonstrated that the application of afatinib (also known as BIBW2992) at reasonable concentrations of around 5 µM leads to structural damage of *in vitro* cultivated metacestode vesicles, induces apoptosis in parasite germinative cells, and when given to infected mice, can lead to a reduction of parasite burden *in vivo* (Cheng et al., 2020). To date, afatinib is therefore the most promising EGFR inhibitor which can potentially be repurposed as a drug against AE. So far, however, it has not been tested which *Echinococcus* EGFR is specifically targeted by afatinib. To clarify this point and to further assess the effects of afatinib on parasite viability, we conducted a series of experiments in which we also included several other EGFR inhibitors that are related to afatinib.

First, we tested the effects of inhibitors on isolated parasite primary cells for cell viability and metacestode vesicle regeneration capacity as well as on mature metacestode vesicles, according to a recently conducted study concerning anti-parasitic activities of PIM kinase inhibitors (Koike et al., 2022). As shown in Figure S1, multiple EGFR inhibitors showed stronger effects against primary cells in the primary screening (Figure S1). We picked up afatinib and dacomitinib (second-generation EGFR receptors), for further analysis, together with osimertinib as a representative of the third generation. As shown in Figure 4A, at concentrations of 1 to 10 µM, afatinib, dacomitinib, and osimertinib significantly affected the viability of isolated primary cells with afatinib showing the most prominent effects. We then also showed that at concentrations of 30 µM and 10 µM, both afatinib and dacomitinib completely inhibited the capacity of primary cells to regenerate metacestode vesicles. With osimertinib, this was only observed at 30 µM. Interestingly, dacomitinib at 3 µM concentration even stimulated the regeneration of metacestode vesicles (Figure 4B). Finally, all three inhibitors also significantly affected the integrity of metacestode vesicles *in vitro*, again with afatinib showing the most prominent effects (Figure 4C). Taken together, these studies verified the negative effects of afatinib on parasite cells and metacestode vesicles as previously observed (Cheng et al., 2020), and introduced dacomitinib and osimertinib as additional EGFR inhibitors with anti-*Echinococcus* activities, albeit being less potent than afatinib.

**Figure 4:**
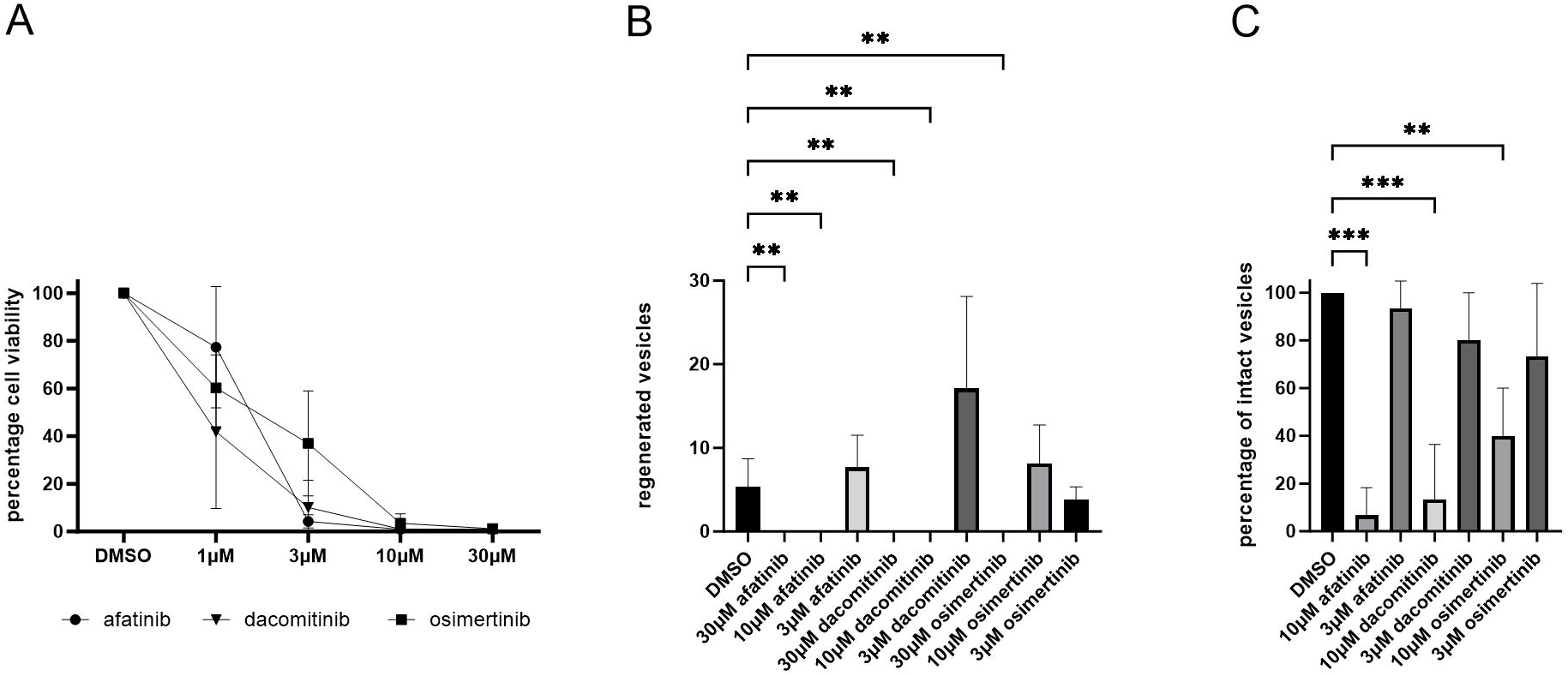
Effects of EGF receptor inhibitors on *Echinococcus* primary cells and metacestode vesicles. (A) Effects of afatinib, dacomitinib, and osimertinib on primary cell culture viability after 3 days of cultivation. Inhibitor concentrations are indicated. (B) Effects of EGF receptor inhibitors on the regeneration of metacestode vesicles from primary cell cultures. Cell cultures were incubated in the presence of inhibitors (concentration as indicated below) for 21 days and regenerated vesicles were counted. (C) Effects of inhibitors on the structural integrity of mature metacestode vesicles after 28 days. Inhibitor concentrations are indicated below. DMSO indicates the negative control. The error bar indicates the standard deviation. P values less than 0.0021 and 0.0002 are summarized with ** and ***.

We were then interested in identifying the target(s) of afatinib. To this end, we made use of the *Xenopus* oocyte system which had previously been used to measure and to demonstrate the ligand induced activities of tyrosine kinase receptor (Leippe et al., 2020) and particularly for *schistosoma mansoni* EGF receptors (Vicogne et al., 2004). In this system, EGFR stimulation by a cognate ligand is required and, according to earlier work (Cheng et al., 2017), we decided to utilize human EGF (HsEGF). Furthermore, we employed the *S. mansoni* EGF receptor SER as a positive control. As shown in Figure 5A, we obtained 30 % GVBD in the case of EmER1 expression and HsEGF stimulation, which is close to what we observed for SER (50 %) and clearly indicates that EmER1 is an active kinase that responds to HsEGF. In the case of EmER2 and EmER3, on the other hand, almost no (EmER2) or no (EmER3) GVBD was obtained, indicating that these receptors are either no enzymatically active kinases or do not functionally interact with HsEGF. We also investigated the level of expression and phosphorylation of all *Echinococcus* EGF receptors in *Xenopus* oocytes and EmER1 as well as EmER2 and EmER3 were well expressed but phosphorylation was only observed for the EmER1 in response to stimulation by HsEGF (Figure S2). This, again, verified that EmER1 is an active kinase that responds to HsEGF whereas EmER2 and EmER3 do not interact with HsEGF.

**Figure 5:**
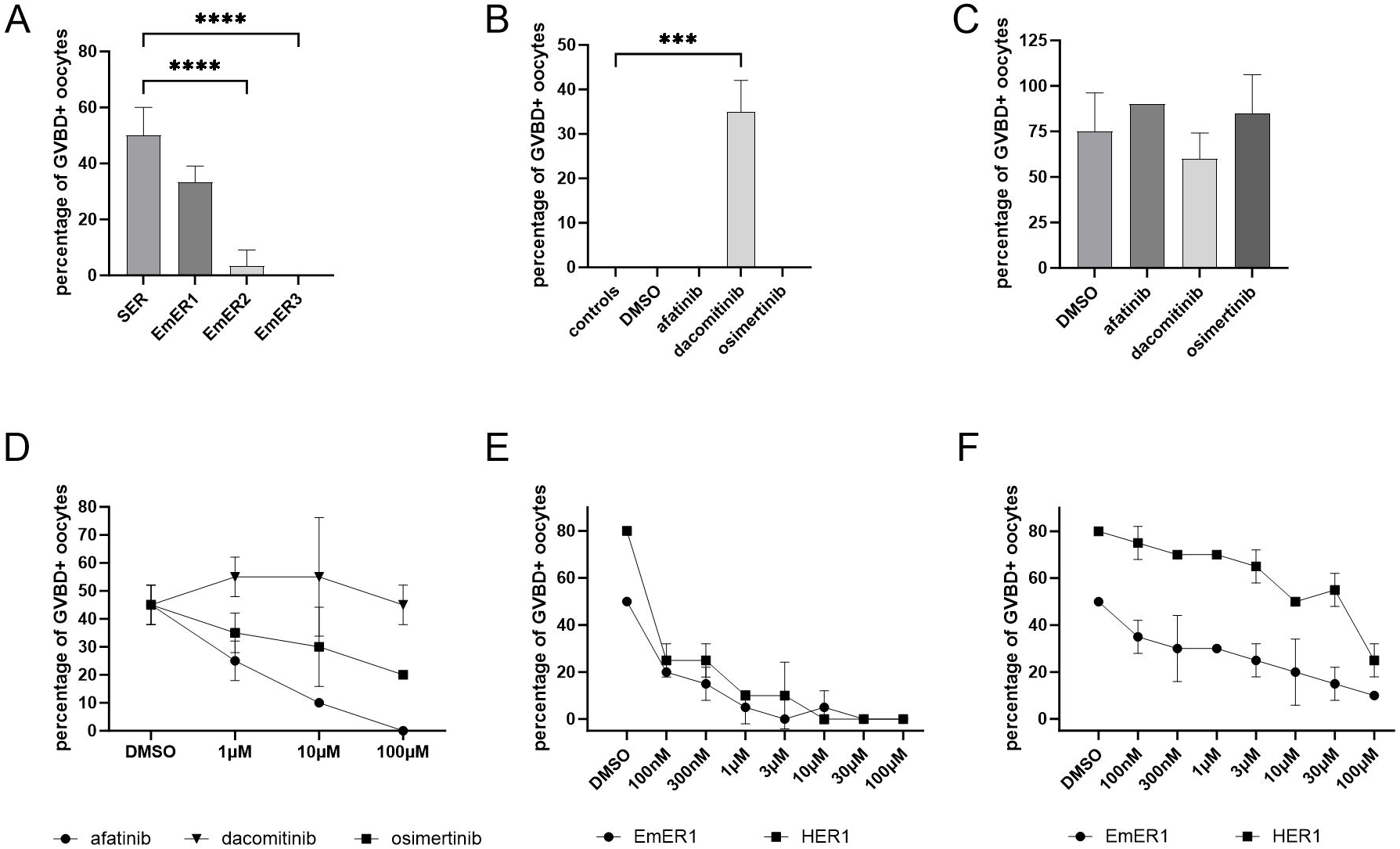
Effects of EGF receptor inhibitors on recombinantly expressed *Echinococcus* receptors. EmER1, EmER2, and EmER3, HER1, as well as the *S. mansoni* receptor SER were expressed in *Xenopus* oocytes and incubated in the presence of human EGF and inhibitors as indicated. Subsequently, germinal vesicle breakdown (GVBD) was measured. (A) Expression of parasite receptors (indicated below) in *Xenopus* oocytes and stimulation by 50 nM human EGF. GVBD was measured after 15 h incubation. 3 technical replicates were used. The error bar represents the standard deviation. p values less than 0.0001 are summarized with ****. (B) *Xenopus* oocytes were incubated in the presence of 10 µM inhibitors for 30 min and GVBD was measured after 15 h incubation. 2 technical replicates were used. The error bar represents the standard deviation. The p values less than 0.0002 are summarized with ***. (C) Percentages of GVBD+ oocytes are shown. Oocytes were treated with 10μM of inhibitors for 30 minutes and stimulated by progesterone. GVBD+ oocytes were counted 15 hours after the stimulation of progesterone. 10 oocytes were used for each condition with 2 technical replicates. Error bars represent standard deviation. (D) Percentages of GVBD+ oocytes are shown. 8 hours after microinjection of EmER1 encoding cRNA, oocytes were treated with 100 nM, 1 μM and 10 μM of inhibitors for 30 minutes and stimulated by 50 nM HsEGF. GVBD+ oocytes were counted 15 h after stimulation by HsEGF. 10 oocytes were used for each condition with 2 technical replicates. Error bars represent standard deviation. (E-F) The percentages of GVBD+ oocytes are shown. 8 hours after cRNA microinjection of EmER1 and HER1, oocytes were treated with 100 nM, 300 nM, 1 μM, 3 μM, 10 μM and 100 μM of afatinib (E) or osimertinib (F) for 30 min and stimulated by human EGF. GVBD+ oocytes were counted 15 hours after EGF stimulation. 10 oocytes were used for each condition with 2 technical replicates. Error bars represent standard deviation.

Having shown that EmER1 is an active kinase we then carried out *Xenopus* oocyte experiments in the presence of EGFR inhibitors. As shown in Figure 5C, *Xenopus* oocytes were well capable of GVBD in response to progesterone stimulation in the presence of afatinib, and osimertinib, indicating that the inhibitors do not have general negative effects on the oocytes, although GVBD was induced by dacomitinib, even without progesterone stimulation (Figure 5B). When we tested these inhibitors on EmER1 expressing *Xenopus* oocytes at concentrations of 1 µM, 10 µM, and 100 µM, we observed a dose-dependent inhibitory effect of afatinib and osimertinib, although the percentage of GVBD with dacomitinib of higher concentration remained high for unknown reason (Figure 5D). Therefore, we then refined our analyses and tested only two inhibitors, afatinib and osimertinib at concentrations from 100 nM to 100 µM (Figure 5E-F). Both inhibitors showed dose-dependent effects. Afatinib was highly effective against both receptors already at 100 nM (Figure 5E). On the other hand, the effect of osimertinib against both receptors was less prominent than that of afatinib at all concentrations, but especially in lower doses, the effect against EmER1 was less prominent against HER1 than EmER1 (Figure 5F).

Taken together, our experiments verified the activities of afatinib on metacestode vesicles previously observed (Cheng et al., 2020) and demonstrated that this compound is also highly effective against isolated parasite cell cultures that are enriched (∼ 80%; Koziol et al., 2014) in germinative cells. Our experiments further showed that EmER1 is an active kinase that is effectively inhibited by afatinib. It is thus highly likely that the effects of afatinib on parasite cells and larvae *in vitro* and on parasite tissue *in vivo* (Cheng et al., 2020) are mostly due to inhibition of EmER1.

### Cloning and characterization of *Echinococcus* EGF ligands

Since asymmetric versus symmetric stem cell division in planarians is regulated by interactions between the *Schmidtea* EGFR and the cognate ligand (Lei et al., 2016) and because it is doubtful whether the *Echinococcus* metacestode during growth *in vivo* has access to sufficiently high concentrations of host EGF for coordinated stem cell regulation, we were interested in identifying possible parasite-intrinsic EGF ligands that might regulate the functions of germinative cells. The hallmark of EGF ligands is the presence of an EGF domain, which contains 6 characteristic Cys residues that form disulfide bridges, but which also occurs in the extracellular regions of numerous other transmembrane proteins (Wouters et al., 2005). In previous bioinformatic analyses, Barberan et al. had applied strict criteria for identifying EGF ligands in metazoans, and thus identified two possible *Echinococcus* EGF ligands (Barberan et al., 2016). One of these, represented by gene model EmuJ_000753300, was assigned to the EGF group of possible ligands, the predicted structure of the second (EmuJ_00090400) suggested that it represents a member of the neuregulin family, which are known as cognate ligands of EGF receptors (Marchionni, 2014). Informed by the *E. multilocularis* genome sequence (Tsai et al., 2013) and recently generated transcriptome data (Herz et al., 2024), we then designed primers to full-length clone and sequence the cDNAs for both genes, which we named *em-egf1* (EmuJ_000753300) and *em-nrg* (EmuJ_000090400), encoding the proteins Em-EGF1 and Em-NRG, respectively.

As shown in Figure 6A, both encoded proteins displayed N-terminal signal peptides and one EGF domain. In the case of Em-NRG, an additional immunoglobulin C-2 domain was present, which is a hallmark of the neuregulin family of EGF ligands (Marchionni, 2014). Furthermore, Em-NRG contained a transmembrane domain, indicating that it is either involved in direct cell-cell communication or that the extracellular portion of the protein must be cleaved to yield a soluble ligand. In both cases, the EGF domain of the putative *Echinococcus* ligands displayed an EGF domain with six characteristic Cys residues that are involved in the formation of disulfide bridges (Figure 6B). We then inspected previous (Tsai et al., 2013) and recent (Herz et al., 2024) transcriptome datasets that we had generated for *E. multilocularis* primary cell cultures and different developmental stages. As shown in Figure 6C, *em-egf1* was mainly expressed in protoscoleces and adult worms, but only showed minor expression in the metacestode. *em-nrg*, on the other hand, showed prominent expression in the metacestode and *em-nrg* transcripts were also enriched after depletion of metacestode vesicles with stem cells, indicating that it is mainly expressed in post-mitotic cells (Figure 6C). Taken together, these analyses indicated that em-nrg displays structural features typical of this family of cytokine ligands and that *em-nrg* is a possible candidate for stem cell regulation in the metacestode stage.

**Figure 6:**
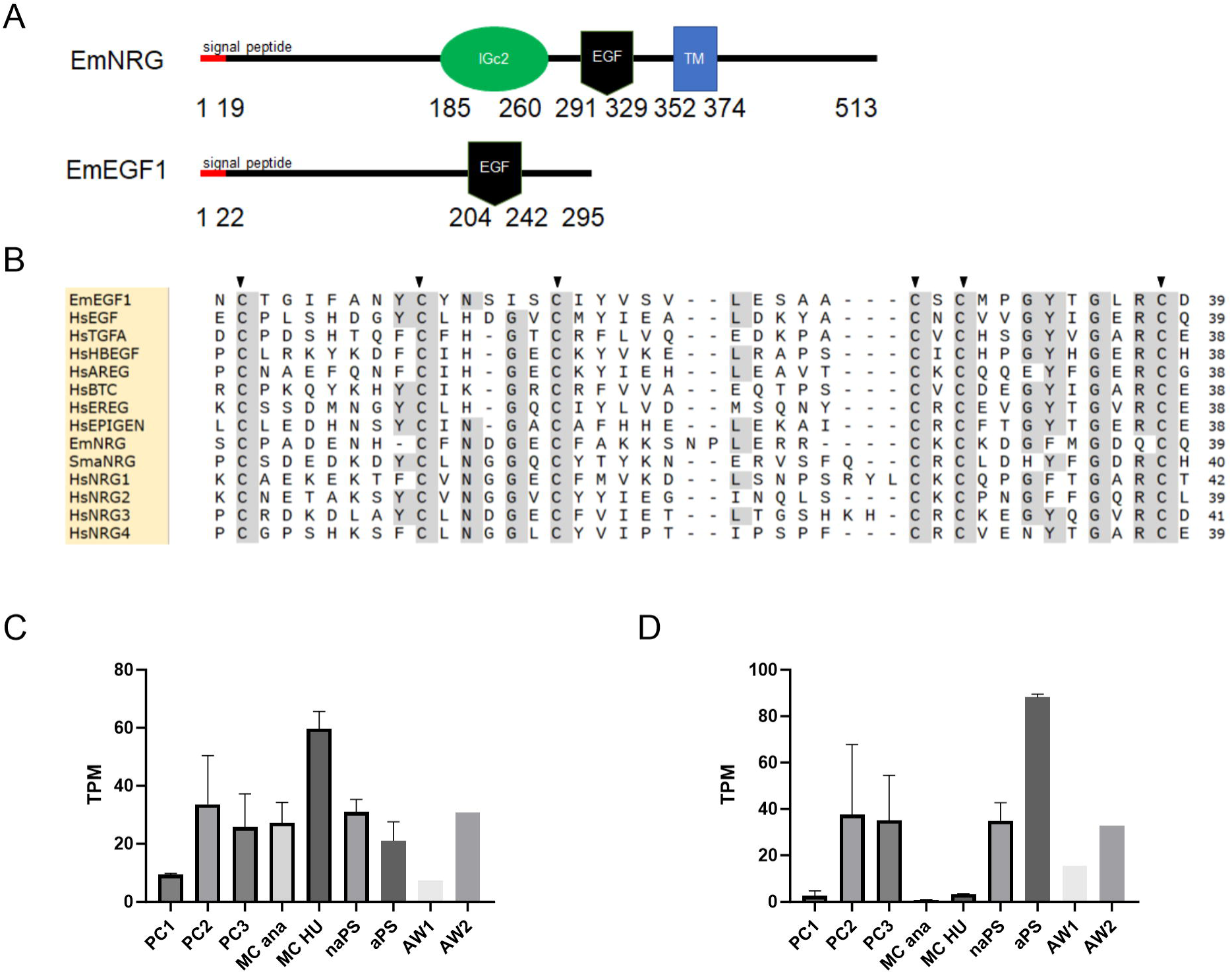
Structure and expression of *Echinococcus* EGF ligands. (A) Domain structure of Em-NRG and Em-EGF1. Displayed are N-terminal signal peptide (red), immunoglobulin type C-2 domain (green), EGF domain (black), and transmembrane region (blue). (B) Amino acid sequence comparison between the EGF domains of EmNRG, EmEGF1, human EGF ligands, and neuregulin of *Schistosoma mansoni* (SmaNRG, Smp_136660). Black triangles indicate 6 characteristic Cys residues that form disulfide bridges. (C-D) Expression of *em-nrg* (C) and *em-egf1* (D) in primary cell cultures at different time points of development (PC1 – PC3; see Figure 2), metacestode vesicles before (MC ana) or after (MC HU) treatment with hydroxyurea, non-activated (naPS) and activated (aPS) protoscolex as well as pre-gravid (AW1) and gravid (AW2) adult worms. Expression values are in TPM and taken from Herz et al. (2024) and Tsai et al. (2013).

### Em-NRG interacts with EmER1 and EmER2

We were then interested in whether the identified *Echinococcus* EGF ligands interact with the three *Echinococcus* EGF receptors. To this end, we employed the MALAR yeast two-hybrid system, which was previously developed to detect extracellular interactions between peptide ligands and cognate receptors (Li et al., 2016). While this system was originally developed to detect interactions between CXC-like chemokine ligands and G-protein coupled receptors (GPCR; Li et al., 2016), it has meanwhile also been employed to detect interactions between neuropeptides and GPCRs (Weth et al., 2020). We now adopted it for measuring the interactions between EGF ligands and possible cognate receptors. For the construction of ligand plasmids, we employed cDNA regions encoding the complete extracellular regions of Em-EGF1, Em-NRG, and human EGF (replacing the N-terminal signal peptide and the transmembrane domains with vector-encoded structures), and for receptor plasmids, the extracellular regions of EmER1, EmER2, EmER3, and the HER1 were used. Unfortunately, Em-EGF1 already induced yeast growth on quadruple dropout plates even with the empty receptor vectors, thus rendering this system unsuitable for measuring EmEGF1 receptor interactions. In the case of Em-NRG, on the other hand, only weaker growth induction was observed with empty receptor vectors.

Since our *Xenopus* oocyte expression experiments had already shown that human EGF stimulates the activities of EmER1 and EmER2 (albeit to much lesser extent), we tested these combinations and, as shown in Figure 7C, in this system the human cytokine interacted with both parasite receptors in a manner comparable to the interaction of human EGF with the cognate human receptor HER1. In the case of EmER3, on the other hand, very weak growth was observed with human EGF. We also tested Em-NRG and observed significant interactions with EmER1, EmER2, and even with human HER1 (Figure 7C; Figure S3). As positive and negative controls we employed previously generated plasmids (Li et al., 2016) for the murine chemokine ligand CXCL12 and the cognate receptor CXCR4 (positive control) as well as CXCL12 with the empty receptor vector (negative control), and, as shown in Figure 7C, they either showed significantly strong or weak growth on quadruple dropout plates. Taken together, these analyses indicated that Em-NRG serves as a ligand for EmER1 and EmER2 in *Echinococcus*.

**Figure 7:**
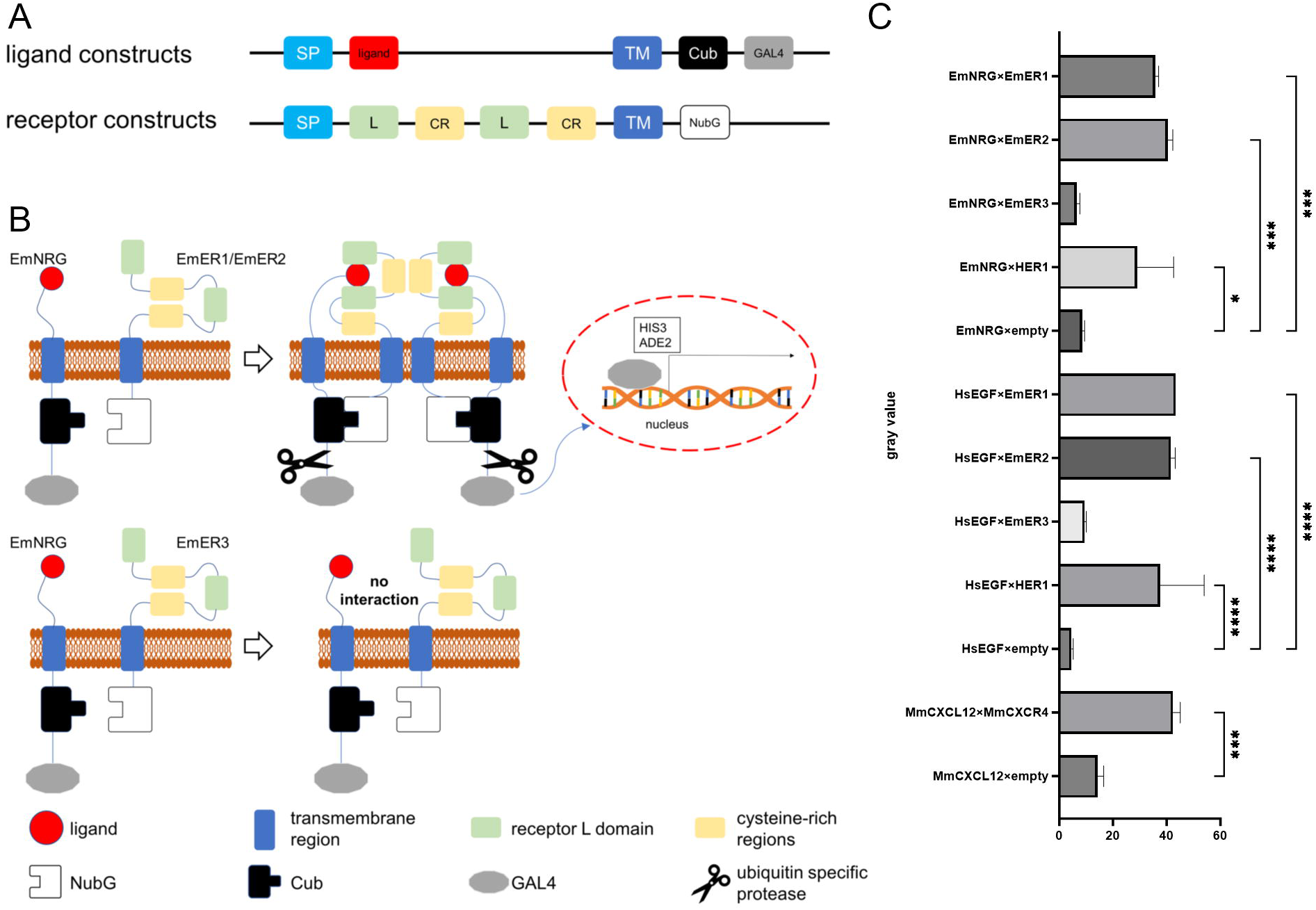
Interactions between *Echinococcus* EGF receptors and ligands. (A) Schematic image of ligand and receptor constructs for MALAR Y2H. SP: host cell signal peptide, L: receptor L domain, CR: cysteine-rich regions, TM: transmembrane region, NubG: N terminal half of ubiquitin with point mutation (Ile13Gly), Cub: C terminal half of ubiquitin. (B) Schematic image of MALAR Y2H principle (modified from Li et al., 2016). (C) The mean level of colony growth quantified as gray value. Error bars represent standard deviation. One-way ANOVA followed by Tukey’s multiple comparison test was used to compare all plasmid combinations to one another, but only the comparisons of corresponding control are shown. P values less than 0.0332, 0.0002, and 0.0001 are summarized with *, ***, and **** respectively. MmCXCL12 and MmCXCR4 are murine chemokine ligands and receptors. They were used as positive control. For images of yeast colonies, see Figure S3.

### *em-nrg* is upregulated during GC clonal expansion

Under steady-state conditions, planarian stem cells are thought to undergo asymmetrical division after which they produce one stem cell (self-renewal) and one differentiating cell (Wagner et al., 2012). We were interested in *em-nrg* expression patterns under comparable conditions and thus performed *em-nrg* specific whole mount *in situ* hybridization (WISH) on *in vitro* cultivated metacestode vesicles combined with EdU incorporation to stain S-phase germinative cells. As shown in Figure 8, we could detect *em-nrg* expression in few germinal layer cells of which the vast majority was not in S-phase. These data indicated that *em-nrg* is either expressed in differentiated cells or differentiating cells.

**Figure 8:**
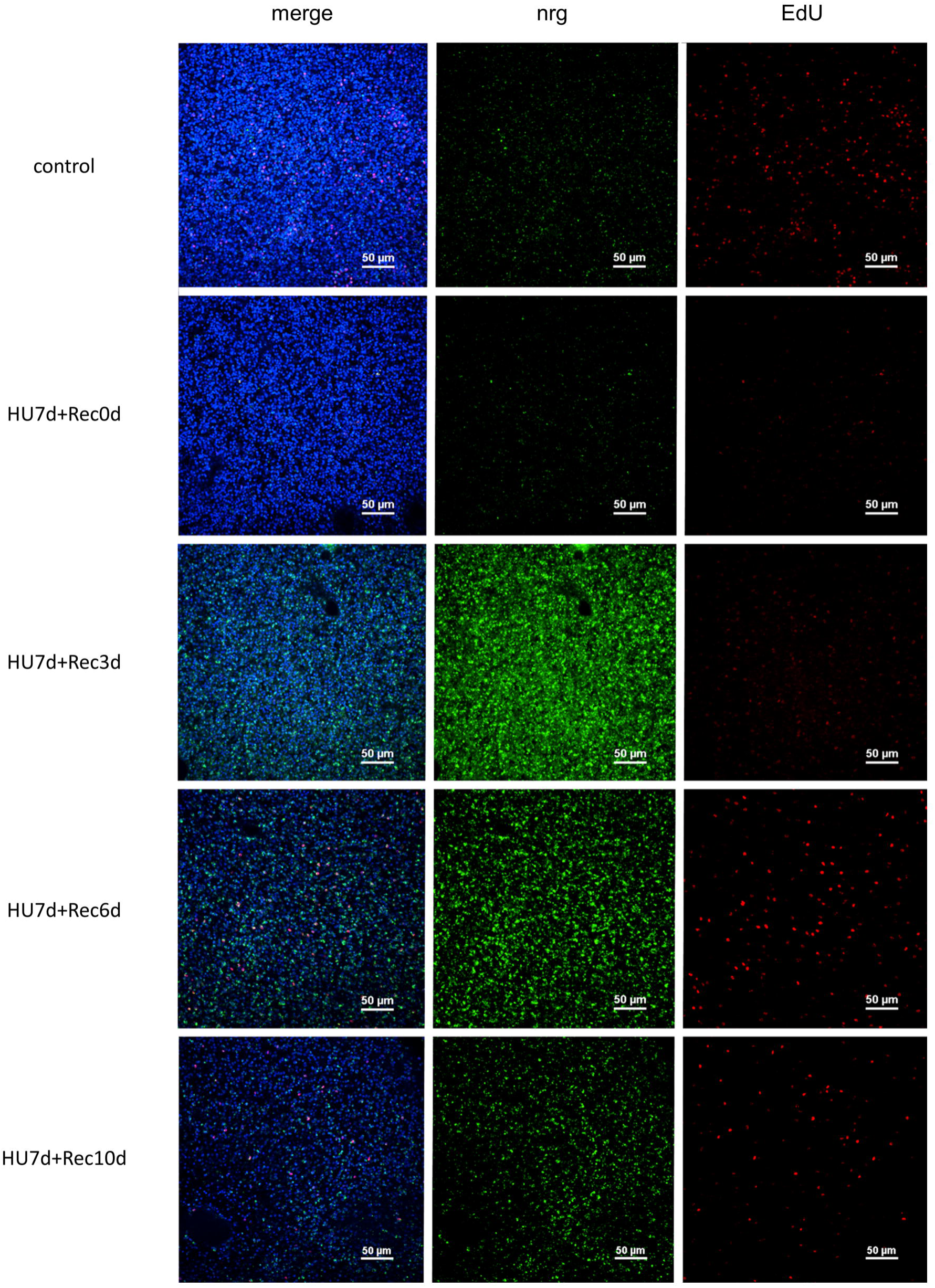
Expression of *em-nrg* during clonal germinal cell expansion. Combined EdU incorporation/WISH assay. Displayed are single confocal slices of the germinal layer of metacestode vesicles after HU treatment and during clonal expansion of stem cells. ‘control’, vesicles without treatment; HU 7d, vesicle after HU treatment for 7 days; Rec, recovery for 3, 6, 10 days as indicated. Channels are green (WISH, em-nrg), red (EdU, S-phase germinative cells, blue (DAPI, nuclei).

Upon diminishing germinative cells in metacestode vesicles by 7 d treatment with hydroxyurea, these undergo several rounds of clonal expansion to replenish the stem cell pool (Koziol et al., 2014). Under these conditions, germinative cells are expected to primarily undergo symmetrical divisions and we were therefore interested in measuring the expression of *em-nrg* after stem cell depletion. To this end, we treated metacestode vesicles with hydroxyurea for 7 d until only few stem cells remained and then allowed the vesicles to recover for 10 d. During this time, we measured the number of EdU+ germinative cells and *em-nrg* expression by WISH. As shown in Figure 7, directly after stem cell depletion only few EdU+ cells were detected and *em-nrg* expression remained low. After 3 d of recovery, we only observed a small increase in EdU+ cells, which agrees with previous studies showing that most stem cells of the metacestode cycle within approximately three days (Koziol et al., 2014; Herz et al., 2024);. The number of cells expressing *em-nrg,* on the other hand, had drastically increased from almost zero to ∼2.250 cells per mm^2^ of metacestode tissue. After 6 d of recovery, the number of EdU+ had sharply increased to ∼500 cells per mm^2^ of metacestode tissue whereas the number of *em-nrg+* cells was slightly reduced (Figure 8). After 10 d recovery, we still observed ∼400 EdU+ cells per mm^2^, whereas the number of *em-nrg+* cells further declined (Figure S4). Taken together, these data indicated that *em-nrg* is particularly highly expressed in metacestode tissue during clonal expansion of stem cells. It is interesting to note that even under conditions of stem cell clonal expansion only very few *em-nrg* expressing cells were found among EdU+ cells (Figure S4), indicating that the gene is mainly expressed in post mitotic cells.

### *em-nrg* is required for metacestode development

We were finally interested in whether RNAi knockdown of *em-nrg* in *E. multilocularis* primary cell cultures influences the regeneration towards metacestode vesicles. To this end, a previously developed protocol for RNAi-based knockdown in primary cell cultures was used (Spiliotis et al., 2010) and, we obtained a knockdown of *em-nrg* expression to around 70% of the GFP- and random controls. Interestingly, both the formation of metacestode vesicles and the viability of primary cell cultures were significantly affected after RNAi against *em-nrg* (Figure 9A-B), indicating that the expression of this EGF ligand encoding gene is necessary for proper stem cell function and parasite development.

**Figure 9:**
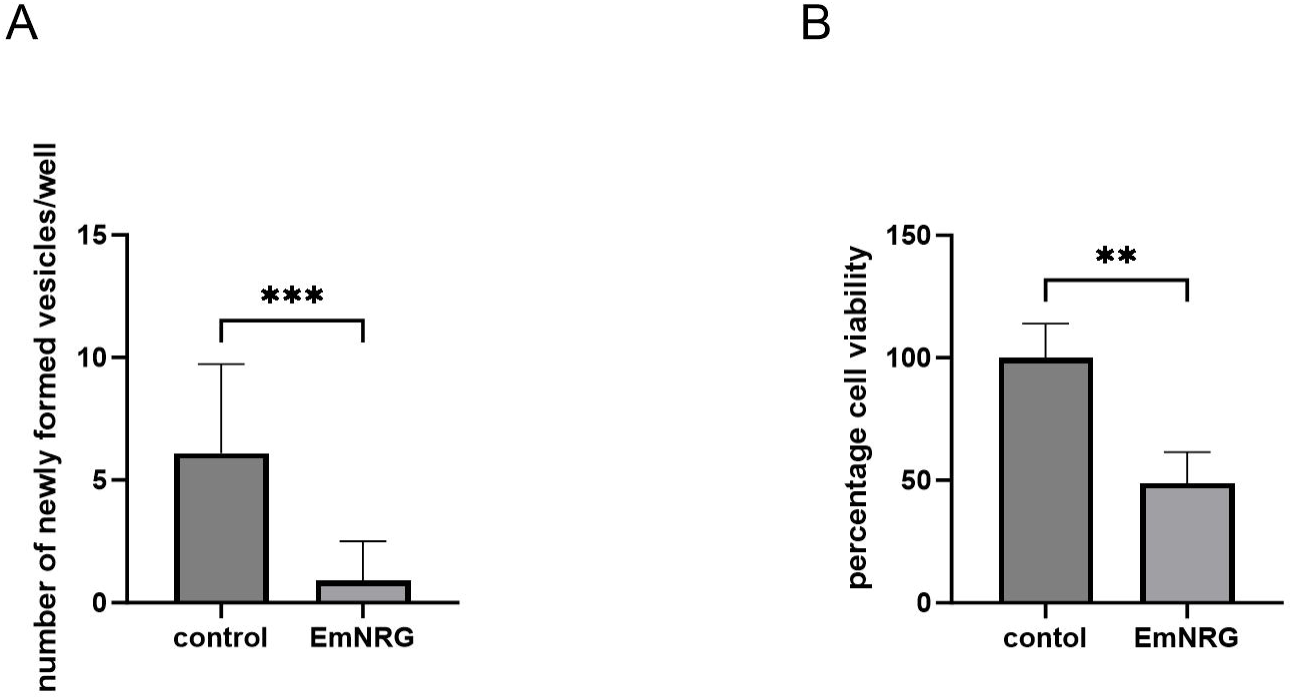
*em-nrg* is required for metacestode vesicle formation. Displayed are the results of RNAi knockdown of *em-nrg*. (A) Formation of mature metacestode vesicles from primary cell cultures after 21 days. The numbers of regenerated vesicles were analyzed by Mann-Whitney test. (B) Cell viability of primary cell cultures after 3 days. The normalized cell viability was analyzed by t-test. P values less than 0.0021, and 0.0002 are summarized with **, and ***, respectively.

## Discussion

The decisive cell type for the growth and proliferation of the *Echinococcus* metacestode within the intermediate host are totipotent parasite stem cells, called germinative cells, which make up 20 – 25 % of all metacestode cells and are the only mitotically active cell type (Brehm and Koziol, 2017). As we have previously shown, under conditions of regular turnover these cells can asymmetrically divide into one stem cell and one differentiating cell (Koziol et al., 2014; Herz et al., 2024). On the other hand, after almost complete stem cell depletion the remaining germinative cells undergo clonal expansion, presumably exclusively through symmetrical divisions (Koziol et al., 2014). The cellular mechanisms governing asymmetrical versus symmetrical division must be tightly controlled and previous research on the related cell type of free-living planarians, the neoblasts, indicated that at least asymmetrical divisions are controlled by EGF signalling (Lei et al., 2016). That EGF signalling also plays an important role in *Echinococcus* stem cell biology has subsequently been supported by data of Cheng et al. who showed that exogenously added human EGF stimulates stem cell proliferation in the *E. multilocularis* metacestode (Cheng et al., 2017) and that the previously characterized EGF receptor EmER1 (Spiliotis et al., 2003) is activated in response to mammalian EGF (Cheng et al., 2017). Based on the data presented herein, we now propose that the mode of *Echinococcus* stem cell division is, at least in part, regulated by the interaction of the newly characterized ligand Em-NRG and EmER1.

According to our structural analyses, Em-NRG belongs to the neuregulin family of EGF molecules that are known to serve as cognate ligands for mammalian EGF receptors and are also involved in stem cell division in planarians (Marchionni, 2014). To characterize Em-NRG receptor interactions, we chose the MALAR yeast two-hybrid system for measuring ligand-receptor interactions and observed a clear interaction between Em-NRG and EmER1 as well as EmER2. We consider this interaction in the yeast two-hybrid system valid since we also observed interactions between human EGF and EmER1 as well as human EGF and the human EGF receptor HER1, both of which have been validated in the *Xenopus* expression system (Figure 5D-F). Furthermore, EmNRG did not interact with EmER3 in the *Xenopus* expression system, and yeast colonies showed very limited growth when it was combined with empty receptor vector. It thus appears that EmER1 and Em-NRG form an interacting receptor-ligand pair in *Echinococcus* EGF signalling. That this interaction involves decisions of symmetrical or asymmetrical division appears plausible since both encoding genes, *emer1* and *em-nrg* are predominantly expressed in post-mitosis when respective decisions are made. To further investigate these aspects, double *in situ* hybridization would have to be performed for *emer1*/*em-nrg* and general stem cell markers such as TRIM (Koziol et al., 2015) or Cip2Ah (Herz et al., 2024), which we have previously described. Furthermore, pulse-chase experiments for specific labeling of stem cell progeny in combination with *emer1*/*emnrg-* specific *in situ* hybridization would be helpful. Interestingly, our *in situ* hybridization experiments indicated that *em-nrg* is strongly upregulated during clonal expansion of germinative cells, in which predominantly symmetric divisions are expected. This contrasts to the ligand-receptor interaction described in planarians by Lei et al, which are involved in asymmetric cell division (Lei et al., 2016). In the planarian system, these questions had been addressed using specific antibodies against EGF receptor and ligand (Lei et al., 2016), and the establishment of respective tools would, of course, also be necessary to investigate questions of symmetric/asymmetric cell division involving EmER1 and EmNRG. In addition, the results of the *Xenopus* expression system and yeast-two-hybrid system do not necessarily indicate that EmER3 is not involved in signal transduction. It is possible that there can be unknown ligands for EmER3, or it might work only when it forms heterodimers with EmER1. In fact, although there are no known ligands for HER2, HER2 works when it dimerizes with other HERs (Kennedy et al., 2016). Because it was impossible to distinguish the outcome of EmER1-EmER3 heterodimer from that of EmER1 homodimer in the *Xenopus* expression assay or MALAR-Y2H, but by improving these methods, this possibility might be verified. Efforts towards such investigations are currently ongoing in our laboratory.

Concerning exploitation of *Echinococcus* EGF signalling for drug development against echinococcosis, both our inhibitor assays and RNAi approaches point to EmER1 as the likely target of afatinib. In the *Xenopus* expression system, EmER1 was clearly inhibited by the drug at concentrations of 100 nM and higher. Furthermore, *Echinococcus* primary cell cultures were drastically affected in their capacity to produce mature vesicles upon RNAi against *emer1*, but not after RNAi against *emer2* or *emer3* (Figure 3). The clear *in vitro* and *in vivo* effects of afatinib previously observed against the *Echinococcus* metacestode (Cheng et al., 2017; Cheng et al., 2020) are therefore likely due to inhibition of EmER1. In this context, it is interesting to note that the amino acid residue that crucially contributes to the activity of afatinib towards mammalian EGF receptors, Cys797, is replaced by Ser in EmER1 (Figure S5A). Afatinib as a second-generation EGFR inhibitor is an aniline-quinazoline derivative that covalently (Figure S5B), and irreversibly, binds to a highly reactive Cys residue within the ATP binding pocket of human EGF receptors, thus suppressing tyrosine kinase activity and preventing receptor dimerization (Nelson et al., 2013). Osimertinib and dacomitinib lead to weaker inhibition of EmER1 in the *Xenopus* expression system and have less prominent effects on cultivated parasite larvae and cells, but the results of respective inhibitor assays performed herein are of relevance to the development of novel drugs against AE. Especially, the third-generation inhibitor osimertinib is applied for the T790M mutation because second-generation inhibitors like dacomitinib and afatinib do not affect receptors with this mutation well, and it has a selective activity to the kinases with this mutation over the wild type (Janne et al., 2015). Although the residue of EmER1 and EmER2 aligned to T790 of HER1 is serine instead of methionine (Figure S5A), in the *Xenopus* expression system, its effect against HER1 at lower doses was weaker than that against EmER1 (Figure 5E-F). In any case, the structural differences between EmER1 and human EGF receptors around the binding site of afatinib could possibly be exploited for the identification of small molecule compounds with high affinity for the parasite receptor molecule, and the comparison of the interaction of osimertinib with HER1 and EmER could result in the identification of lead compounds with selective affinity. A feasible approach towards this end would be AI-based *in silico* pre-screening of very large compound libraries similar to what we recently conducted for the *Echinococcus* PIM kinase (Koike et al., 2022), followed by biochemical assays on parasite cell cultures and EmER1, recombinantly expressed in the *Xenopus* expression system. Current systems of AI-based *in silico* screening of compound libraries strongly rely on the information on differential binding affinities of diverse compounds towards the target molecule (Kim et al., 2020).

## Conclusions

We herein identified the *E. multilocularis* receptor tyrosine kinase EmER1 as a target for afatinib, which has been shown to exert prominent anti-parasitic activities both *in vitro* and *in vivo*. Together with our studies on related compounds such as osimertinib and dacomitinib, these data are highly relevant for the development of novel anti-infectives that target EmER1, e.g. by AI-based screening of compound libraries, followed by inhibitor assays in the *Xenopus* system. We also identified a parasite-derived, cognate ligand for EmER1, EmNRG, that is drastically upregulated during clonal expansion of *Echinococcus* germinative cells and that is required for the formation of parasite vesicles from parasite stem cell cultures. Although we cannot exclude that host EGF, which is induced during regenerative processes of the liver, influences parasite proliferation during an infection, we propose that EmNRG is the most important ligand for EmER1, and that this cognate receptor-ligand pair regulates asymmetric/symmetric division decisions of germinative cells during parasite development. These data are highly relevant for elucidating the differentiation mechanisms of *Echinococcus* germinative cells, which are the most important cell type for parasite growth and establishment during AE.

## Acknowledgements

This work was supported by a grant of the Bayerische Forschungsstiftung [AZ-1341-18] (to KB and SH). AK was supported by a fellowship of the Heiwa Nakajima Foundation under grant number [AK-1-2019]. The authors acknowledge technical assistance from Dirk Radloff and Monika Bergmann. We also wish to thank researchers in Han laboratory (Shanghai Jiao Tong university), Christoph Grevelding and Oliver Weth for helpful suggestions and for providing us with MALAR Y2H plasmids.

## Supplimentary Figure Legends

**Supplimentary Figure S1: Activities of protein kinase inhibitors against primary cells of *E. multilocularis*** (A-B) Heatmap showing the effects of kinase inhibitors on primary cells of *E. multilocularis*. The color code below indicates the percentage of luminescence signal (proportional to the number of viable cells), normalized to signals from DMSO controls, after 3 days of incubation with 3-30 μM of inhibitor. Inhibitor names, human target proteins, and kinase groups are indicated in the table to the left. (A) is the result of the screening with the second inhibitor list and (B) is that with the third inhibitor list. The results with the first inhibitor list were previously published (Koike et al, 2022).

**Supplimentary Figure S2; Phosphorylation of the recombinant protein** 24 hours after the microinjection of cRNA or not (-), the oocytes were stimulated by human EGF for 15 hours. The recombinant protein expressed in the oocytes was purified with myc-IP from the lysates and phosphorylation of the recombinant protein was evaluated by western blot with PY20 antibody.

**Supplimentary Figure S3: Yeast growth on the selective plates** Representative images of yeast transformants grown on double dropout (-Leu/-Trp, selective for plasmids), triple dropout (-Leu/-Trp/-His, indicating interactions under moderate stringency), and quadruple dropout plates (-Leu/-Trp/-His/-Ade, indicating interactions under high stringency). Inoculation densities are shown above, and plasmid combinations are shown on the left. The raw images were converted into grayscale and the background was subtracted with the algorism of sliding paraboloid.

**Supplementary Figure S4: WISH signal of *em-nrg* during clonal germinal cell expansion.** (A-B) The mean number of cells with each signal (per mm^2^). (A) *in situ* signal+ cells, (B) EdU+ cells. The error bars represent the standard deviation. One-way ANOVA followed by Dunnett’s multiple comparison tests was used to compare the control and each treatment condition. P values less than 0.0332, 0.0002, and 0.0001 are summarized with *, ***, and **** respectively.

**Supplementary Figure S5**: **Important residues for the interaction between EGF receptors and afatinib** (A) Part of the amino acid sequence alignment of HER1, HER3, and EmER1-3. The N terminal part of the kinase domain was excised. Residues identical to HER1 are shown in gray highlight. Black triangles indicate residues of HER1 interacting with afatinib (numbers are positions on HER1). Thr790 and Cys797, which affect the toxicity of EGFR inhibitors, are shown with blue letters. The red triangle indicates Asparagine 834 of HER3 (number here is the position on HER3), which makes it catalytically impaired. Hinge region, DFG motif, and E746-A750, residues frequently lost in exon 19 deletion mutant, are also shown. (B) Residues of wild-type HER1 interacting with afatinib are shown in yellow letters and especially important residues are shown in purple on the 3D model. The white arrow indicates the covalent bond at Cys797. This image was visualized with Molmil (Protein Data Bank Japan) from the crystal structure data deposited to Protein Deta Bank (ID: 4G5J).

(C) Residues of wild-type HER1 interacting with afatinib and aligned residues in other receptor kinases. Residues identical to HER1 are shown in yellow highlight, and those similar to HER1 are shown in gray. The percentage of identical and similar residues is shown in the lowest two rows.

## Supplementary Table Information

**Table S1**: Protein kinase inhibitors

**Table S2**: primers

**Table S3**: accession numbers

**Table S4**: siRNA sequences

