## Supplementary figures and images for "Putative EGF ligand and receptor of *Echinococcus multilocularis* that are critical for parasite development"

### Supplemental Figure S1

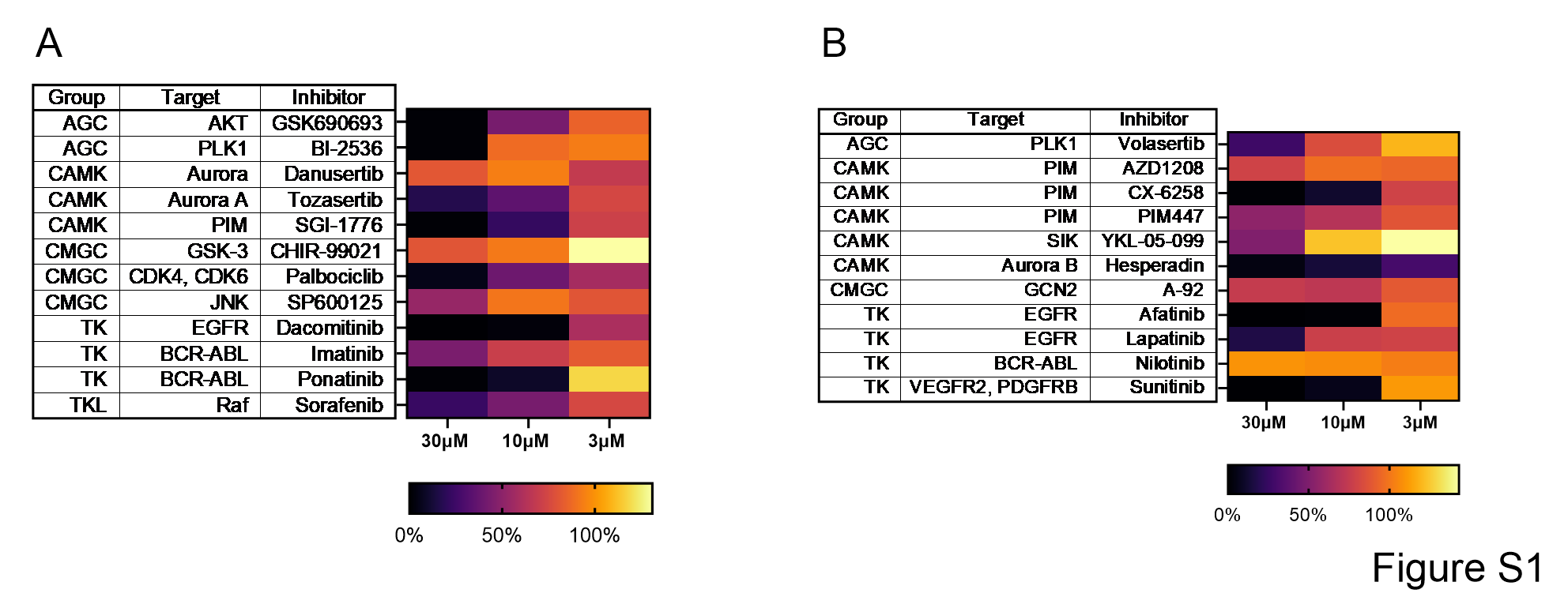

### Supplemental Figure S2

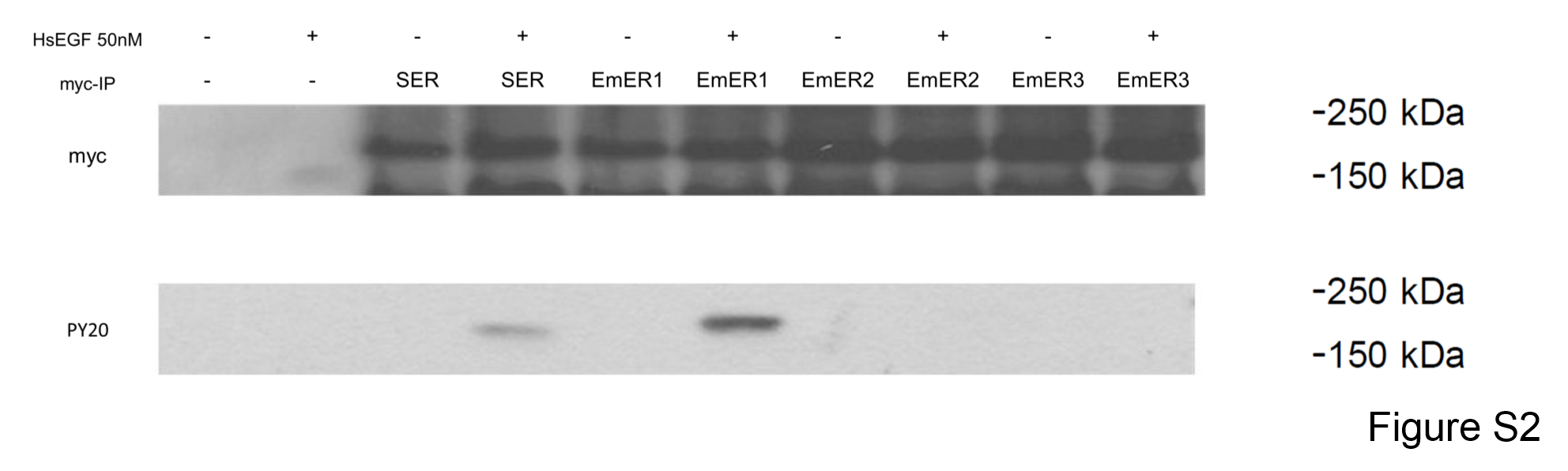

### Supplemental Figure S3

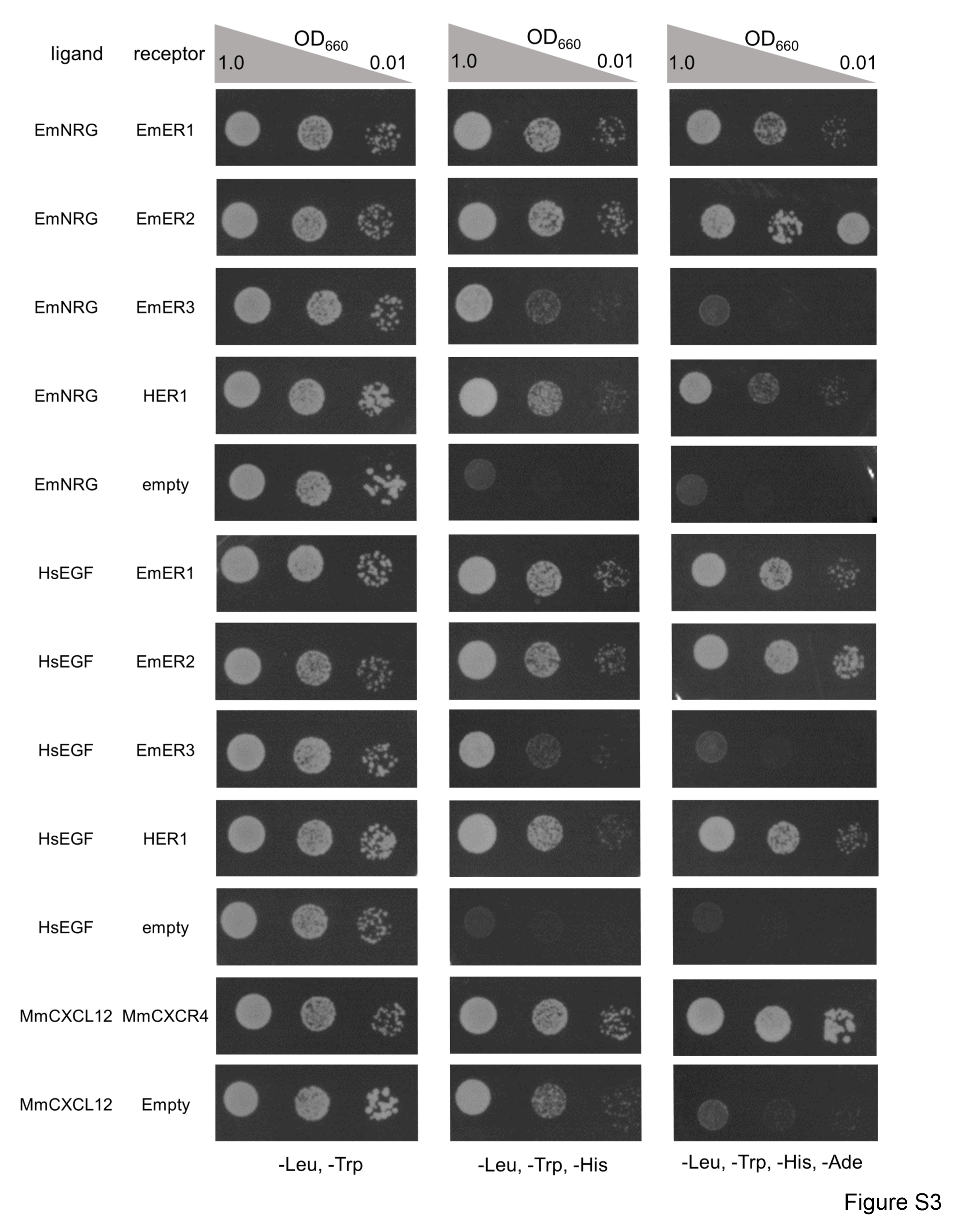

### Supplemental Figure S4

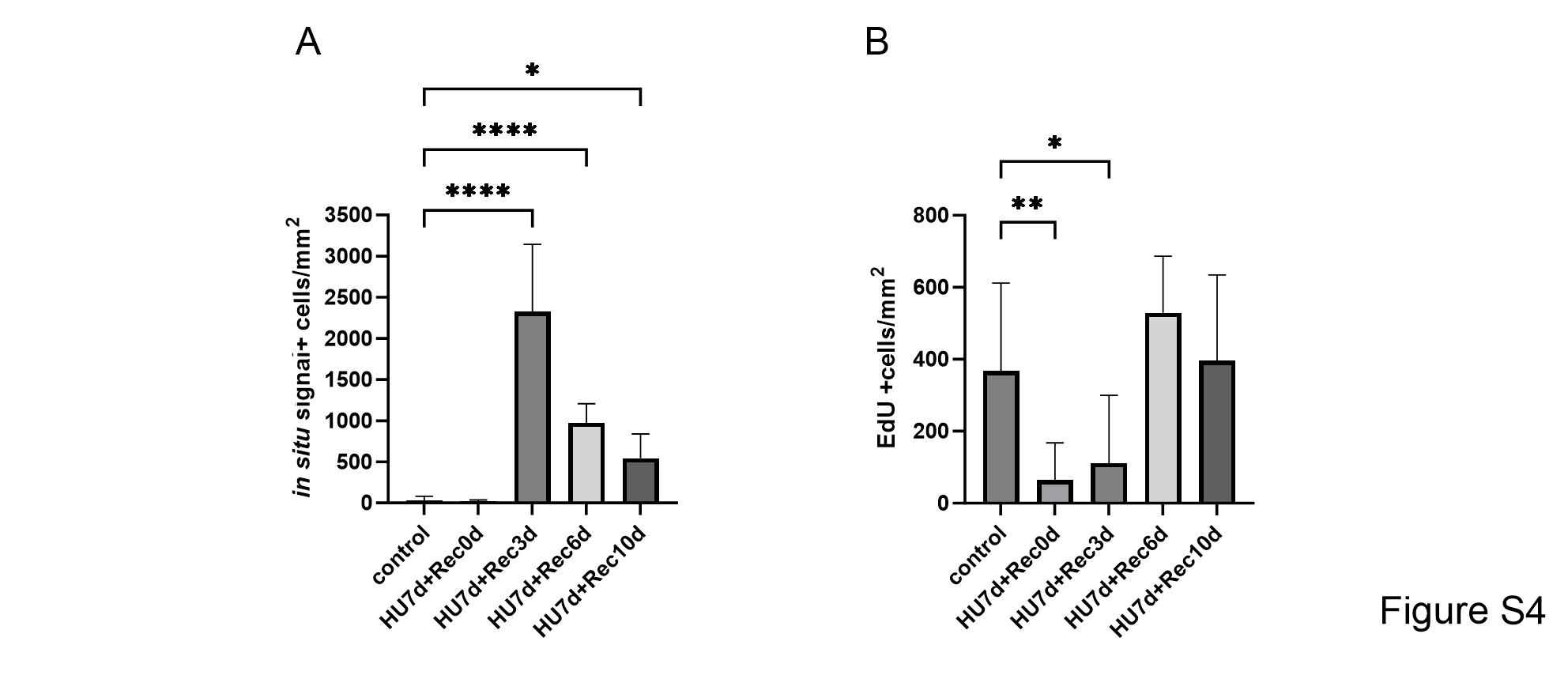

### Supplemental Figure S5

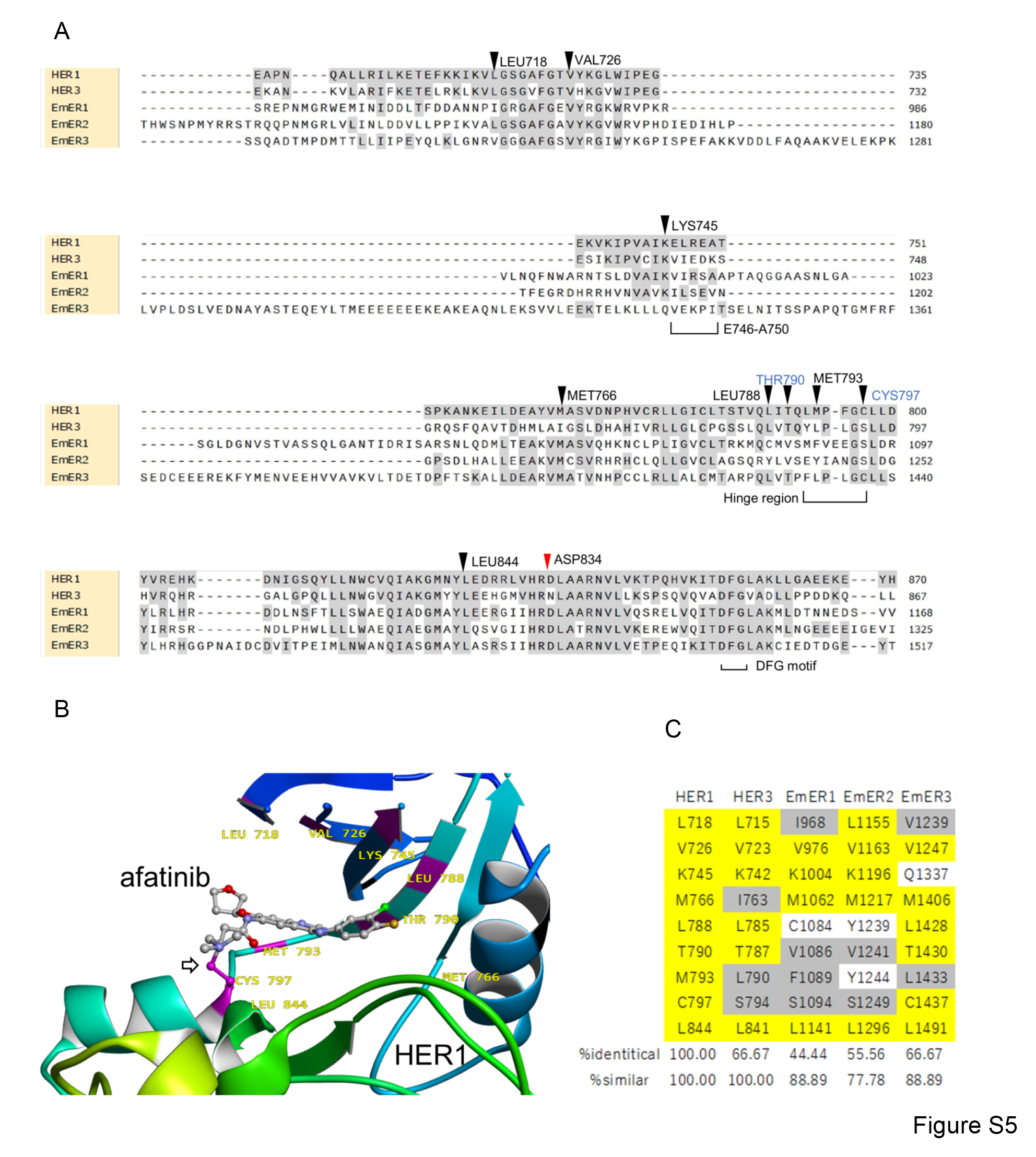
