## Supplemental Table S1 for "Putative EGF ligand and receptor of *Echinococcus multilocularis* that are critical for parasite development"

Second set of screening (Fig. S1A)

| Kinase inhibitor | Provider |
|---|---|
| CHIR-99021 | Selleckchem |
| dacomitinib | abcr GmbH |
| danusertib | abcr GmbH |
| GSK690693 | Tocris |
| imatinib | Sigma-aldrich |
| palbociclib | Selleckchem |
| ponatinib | abcr GmbH |
| SGI-1776 | Selleckchem |
| sorafenib | abcr GmbH |
| SP600125 | Selleckchem |
| Bi2536 | Axon Medchem |
| tozasertib | Selleckchem |

Third set of screening (Fig. S1B)

| Kinase inhibitor | Provider |
|---|---|
| A-92 | Axon Medchem |
| AZD1208 | Selleckchem |
| afatinib | Cayman chemical |
| CX-6258 | Cayman chemical |
| hesperadin | Cayman chemical |
| lapatinib | Cayman chemical |
| nilotinib | Selleckchem |
| PIM447 | Selleckchem |
| sunitinib | Hölzel Diagnostika Handels |
| volasertib | Cayman chemical |
| YKL-05-099 | Hölzel Diagnostika Handels |

Other

| Kinase inhibitor | Provider |
|---|---|
| AZD9291 (osimertinib) | Cayman chemical |

Table S1
