## Supplemental Table S2 for "Putative EGF ligand and receptor of *Echinococcus multilocularis* that are critical for parasite development"

| plasmid name | primer name | 5'-sequence-3' |
|---|---|---|
| pCR™-XL-2-TOPO | CD3-RT | ATC TCT TGA AAG GAT CCT GCA GGT |
| pSec-Tag2/Hygro in general | pSTH_Seq_Fw | GCTCGAGGAGGGCCCGAA |
| pSec-Tag2/Hygro in general | pSTH_Seq_Rev | CTGGGCCGCGTCACCAGT |
| pSec-Tag2/Hygro-EmER1 | STHEmER1-Fwd | CCACTGGTGACGCGGCCCAGTTTAAGCTATCCCTTGCCCC |
| pSec-Tag2/Hygro-EmER1 | STHEmER1-Rev | TGTTCGGGCCCTCCTCGAGCGTAGAAATGAAGTTGCTTGGCGC |
| pSec-Tag2/Hygro-EmER1 | EmER1seq3529Rev | TCCAGACATCGGTTTTGTGA |
| pSec-Tag2/Hygro-EmER2 | STHEmER2-Fwd | CCACTGGTGACGCGGCCCAGTTTGTAACAGATGTAGGAAAATATC |
| pSec-Tag2/Hygro-EmER2 | STHEmER2-Rev | TGTTCGGGCCCTCCTCGAGCCAAGTCAGTGGCATCAGTAGA |
| pSec-Tag2/Hygro-EmER2 | EmER2seq3357Rev | CCAGTGAGTGCTGCGATAAA |
| pSec-Tag2/Hygro-EmER3 | STHEmER3-Fwd | CCACTGGTGACGCGGCCCAGGCTGAAGATTTCCCTACCATTG |
| pSec-Tag2/Hygro-EmER3 | STHEmER3-Rev | TGTTCGGGCCCTCCTCGAGCAGCAACCTCCTCTGGTTCAGC |
| pSec-Tag2/Hygro-EmER3 | EmER3seq3753Rev | CTGCGCAAATAAGTCGTCAA |
| pSec-Tag2/Hygro-EmER1/pGBKT7_SP_OST1_EmER1_NubG | EmER1seq1590Fwd | CTGTGTCTCCGATTGCTCTG |
| pSec-Tag2/Hygro-EmER1/pGBKT7_SP_OST1_EmER1_NubG | EmER1cpRev1776Rev | GCATTTGTGCCACTTGTTTCATTGAAAGTGGC |
| pSec-Tag2/Hygro-EmER2/pGBKT7_SP_OST1_EmER2_NubG | EmER2seq1530Fwd | TCGTCGTGAAGATTGTCTGC |
| pSec-Tag2/Hygro-EmER2/pGBKT7_SP_OST1_EmER2_NubG | EmER2cp1735Rev | GCGGTCATGGGGTTGCTCTGAGGTGGCAATG |
| pSec-Tag2/Hygro-EmER2/pGBKT7_SP_OST1_EmER2_NubG | EmER3seq1256Fwd | GCTTGGCCTGCAGAAGATAC |
| pSec-Tag2/Hygro-EmER3/pGBKT7_SP_OST1_EmER3_NubG | EmER3cp1810Rev | TTGTCCGTTCCAACATAGAGAACACCCGTTATTTC |
| pGAD SP-WBP1_cloning_linker_TMP_Cub_ GAL4 | LigSeqFwd | ccaagcatacaatcaactccaa |
| pGAD SP-WBP1_cloning_linker_TMP_Cub_ GAL4 | LigSeqRev | gcggtctggaaggatgaat |
| pGAD SP-WBP1_EmEGF_linker_TMP_Cub_ GAL4 | LigEmEGF-Fwd | gcaggtgtacttcaactcccTCTACGATCCGGCCGCGT |
| pGAD SP-WBP1_EmEGF_linker_TMP_Cub_ GAL4 | LigEmEGF-Rev | ccggtcgtaccagatcccccAACTTCGTCCTCTATTTCGAAGGCC |
| pGAD SP-WBP1_EmNRG_linker_TMP_Cub_ GAL4 | LigEmNRG-Fwd | gcaggtgtacttcaactcccGCCATGGAATCCAGAGATTTG |
| pGAD SP-WBP1_EmNRG_linker_TMP_Cub_ GAL4 | LigEmNRG-Rev | ccggtcgtaccagatcccccTTGCAACCAAATGGTATTCG |
| pGAD SP-WBP1_HsEGF_linker_TMP_Cub_ GAL4 | LigHsEGF-Fwd | gcaggtgtacttcaactcccAATAGTGACTCTGAATGTCCCCTG |
| pGAD SP-WBP1_HsEGF_linker_TMP_Cub_ GAL4 | LigHsEGF-Rev | ccggtcgtaccagatcccccGCGCAGTTCCCACCACTTC |
| pGBKT7_SP_OST1_cloning_NubG | RecepSeqFwd | cttgaagcaagcctcctgaa |
| pGBKT7_SP_OST1_cloning_NubG | RecepSeqRev | cattccttgtcttgaattttcg |
| pGBKT7_SP_OST1_EmER1_NubG | RecEmER1-Fwd | agcccatggcttcaactcccTTTAAGCTATCCCTTGCC |
| pGBKT7_SP_OST1_EmER1_NubG | RecEmER1-Rev | gatccaccttctaggatcccGCGTTTGGCCTTGTAATTG |
| pGBKT7_SP_OST1_EmER2_NubG | RecEmER2-Fwd | agcccatggcttcaactcccTTTGTAACAGATGTAGGAAAATATCGAG |
| pGBKT7_SP_OST1_EmER2_NubG | RecEmER2-Rev | gatccaccttctaggatcccTTTGTGGCGCCCTTTGTAG |
| pGBKT7_SP_OST1_EmER3_NubG | RecEmER3-Fwd | agcccatggcttcaactcccGCTGAAGATTTCCCTACC |
| pGBKT7_SP_OST1_EmER3_NubG | RecEmER3-Rev | gatccaccttctaggatcccAATTACAATGCGAGAGCAC |
| pGBKT7_SP_OST1_HER1_NubG | RecHER1-Fwd | agcccatggcttcaactcccCTGGAAGAGAAGAAAGTGTGCC |
| pGBKT7_SP_OST1_HER1_NubG | RecHER1-Rev | gatccaccttctaggatcccCACGATGTGTCTCCGCCG |
| pGBKT7_SP_OST1_HER1_NubG | HER1Seq817Fwd | CGGAACTACGTGGTCACAGA |
| pJet1.2 based in general | T7 Plus2 | AGA AGA GTA ATA CGA CTC ACT ATA GG |
| pJet1.2 based in general | 5-Sp6+pJet1.2 Rev-3 | ATA ATT TAG GTG ACACTA TAG AAC ATC GAT TTT CCA TGG CAG |
| pJet-EmER1 | EmER1insituFwd | GTTCCCTGTGCAAAAACTC |
| pJet-EmER1 | EmER1insituRev | CAGAGACGCTAAACACTTGG |
| pJet-EmER2 | EmER2insituFwd | CACTTGCAGCCTCTTTTC |
| pJet-EmER2 | EmER2insituRev | CTTCGTCTCTATGGTCGG |
| pJet-EmER3 | EmER3insituFwd | cgccaccttattcaatatc |
| pJet-EmER3 | EmER3insituRev | ggtgttgtacttgtgaaacttc |
| pJet-EmNRG_fulllength | EmNRGfulllengthFwd | ATGATGGCTATATTTGTATTAGTC |
| pJet-EmNRG_fulllength | EmNRGfulllengthRev | CTACAAGAAATTGGGATTTTC |
| pJet-EmNRG_insitu | EmNRGinsituFwd | TCGTACCAGTGGAATCGTTTT |
| pJet-EmNRG_insitu | EmNRGinsituRev | CGTCTACCTCCTGCTCTTCG |

Table S2
