## Supplemental Table S3 for "Putative EGF ligand and receptor of *Echinococcus multilocularis* that are critical for parasite development"

Human receptors

| Protein | UniProt ID |
| --- | --- |
| HER1 | P00533 |
| HER2 | P04626 |
| HER3 | P21860 |
| HER4 | Q15303 |

Human ligands

| Protein | UniProt ID |
| --- | --- |
| HsNRG1 | Q02297 |
| HsNRG2 | O14511 |
| HsNRG3 | P56975 |
| HsNRG4 | Q8WWG1 |
| HsEGF | P01133 |
| HsTGFA | P01135 |
| HsHBEGF | Q99075 |
| HsAREG | P15514 |
| HsBTC | P35070 |
| HsEREG | O14944 |
| HsEPGN | Q6UW88 |

Platyhelminth receptors

| species | protein | UniProt ID | WormBase ParaSite ID | GenBANK ID |
| --- | --- | --- | --- | --- |
| *Echinococcus multilocularis* | EmER1 | A0A087VXU7 | EmuJ_000075800 | AJ515524 |
| *Echinococcus multilocularis* | EmER2 | A0A0S4MP42 | EmuJ_000617300 | PP616682 |
| *Echinococcus multilocularis* | EmER3 | A0A068YB41 | EmuJ_000969600 | PP616683 |
| *Schistosoma mansoni* | SER | A0A3Q0KJB8 | Smp_093930 | |
| *Schistosoma mansoni* | | A0A3Q0KS13 | Smp_165470 | |
| *Schistosoma mansoni* | | A0A5K4FD90 | Smp_344500 | |
| *Schmidtea mediterranea* | | A0A1B1ACX9 | h1SMcG0020891 | |
| *Schmidtea mediterranea* | | A0A1B1ACY2 | h2SMcG0000358 | |
| *Schmidtea mediterranea* | | A0A1B1ACY3 | h1SMcG0011666 | |
| *Schmidtea mediterranea* | | A0A1B1ACY5 | h2SMcG0014530 | |
| *Schmidtea mediterranea* | | A0A1B1ACY9 | h2SMcG0001820 | |

Platyhelminth ligands

| species | protein | UniProt ID | WormBase ParaSite ID | GenBANK ID |
| --- | --- | --- | --- | --- |
| *Echinococcus multilocularis* | EmNRG | A0A087VYF8 | EmuJ_000090400 | PP616684 |
| *Echinococcus multilocularis* | EmEGF1 | A0A068Y670 | EmuJ_000753300 | PQ757332 |
| *Schistosoma mansoni* | SmaNRG | A0A3Q0KMS8 | Smp_136660 | |
| *Schmidtea mediterranea* | SmeNRG | A0A172MAV5 | h1SMcG0014188 | |
| *Schmidtea mediterranea* | | A0A161HX36 | h2SMcG0016877 | |
| *Schmidtea mediterranea* | | A0A1B1ACX5 | h1SMcG0022115 | |
| *Schmidtea mediterranea* | | A0A1B1ACX7 | h1SMcG0016670 | |
| *Schmidtea mediterranea* | | A0A1B1ACX8 | h2SMcG0000456 | |
| *Schmidtea mediterranea* | | A0A1B1ACY0 | mk4.000428.00 | |
| *Schmidtea mediterranea* | | A0A1B1ACY1 | h2SMcG0007982 | |

Table S3
